# Four SpsP neurons are an integrating sleep regulation hub in *Drosophila*

**DOI:** 10.1101/2024.09.02.610881

**Authors:** Xihuimin Dai, Jasmine Quynh Le, Dingbang Ma, Michael Rosbash

**Affiliations:** Howard Hughes Medical Institute, Brandeis University, Waltham, MA 02454; Interdisciplinary Research Center on Biology and Chemistry, Shanghai Institute of Organic Chemistry, Chinese Academy of Sciences, Shanghai, China

## Abstract

Sleep is an essential and conserved behavior, yet the mechanisms underlying sleep regulation remain largely unknown. To address the neural mechanisms of sleep drive, here we carry out whole brain calcium-modulated photoactivatable ratiometric integrator (CaMPARI) imaging of *Drosophila* and show that the activity of the protocerebral bridge (PB), a part of the central complex, correlates with sleep drive. Through a neural activation screen followed by anatomical and functional connectivity assays, we further narrow down the key player of sleep regulation in the PB to a three-layer circuit composed of 4 SpsP neurons and their upstream and downstream synaptic partners: the 4 SpsP neurons act as an integrating hub by responding to ellipsoid body (EB) signals from EPG neurons, and by sending signals back to the EB through PEcG neurons. Moreover, sleep deprivation enriches the presynaptic active zones of SpsP neurons and strengthens the connections of the EPG-SpsP-PEcG circuit, indicating plasticity gating in the circuit in response to sleep drive change. As the SpsP neurons also receive input from the sensorimotor brain region and given their known role in navigation, these neurons potentially further integrate sleep drive with other sensorimotor cues. The data taken together indicate that the four SpsP neurons and their sleep regulatory circuit play an important and dynamic role in sleep regulation.

## Introduction

Sleep is important for multiple biological processes, which include but are not limited to immune responses, learning and memory, and waste clearance^1^. While sleep is often defined by behavioral changes such as prolonged immobility, reduced responsiveness to external stimuli, and sleep recovery after deprivation^2,3^, sleep stages and wakefulness are traditionally distinguished by the electrophysiological state of the brain. Indeed, mammalian sleep stages and wakefulness are characterized by distinct electroencephalographic (EEG) patterns, and different sleep stages have even been identified in *Drosophila*, in this case through local field potential (LFP) recordings^4–7^ and Hidden Markov Modeling of sleep behavior^8^. This concordance suggests that sleep is regulated by discrete brain neural circuits in flies as well as mammals and suggests that some fly neurons should be more active when the animal needs to sleep or needs to maintain sleep as has been shown in mammals^9^.

This search for sleep-relevant neurons overlaps with a basic principle of sleep regulation, namely, the importance of a circadian as well as a homeostasis process^10^. The circadian process is controlled by the ∼150 circadian clock neurons in *Drosophila*^11^, but fly “sleep centers” remain poorly defined, especially those that change with sleep state, deep versus light sleep for example, or brain regions that are critical for sleep homeostasis.

Recent studies have suggested that the central complex, especially its dorsal fan-shaped body (dFB) and EB neurons, are important for the regulation of sleep and sleep homeostasis. Manipulation of drivers labeling the dFB neurons or the EB neurons caused changes in sleep or sleep homeostasis, and functional studies proposed a recurrent circuit between the dFB neurons, the helicon cells and the EB neurons, which incorporates signals from the circadian neurons^12,13^. Nonetheless, these studies have been constrained by the modest availability and specificity of individual Gal4 drivers, and have limited insights into dynamic neuronal activity patterns across the fly brain during natural changes in sleep drive. On the other hand, a recent study has revealed that distinct neuronal ensembles are active during different sleep states^14^, but the identities and projection patterns of these active neurons remain unknown.

To overcome this limitation of the Gal4 system and search for neurons that respond to sleep drive change in a natural context, we instead began by using CaMPARI to take an irreversible snapshot of whole brain calcium activity in sleep-deprived flies. The results showed that the protocerebral bridge (PB), another component of the central complex, exhibits a substantially higher calcium activity in flies with high sleep drive than any other easily detectable brain location. Following a thermogenetic activation screen with sparsely labeled split Gal4 drivers, we identified 3 sleep-promoting subsets of PB neurons, namely, 4 SpsP neurons, ∼20 pairs of EPG neurons, and ∼9 pairs of PEcG neurons. The 4 SpsP neurons alone are strongly sleep promoting and show consistent inhibition as well as activation sleep phenotypes. Through connectivity assays, we further show that they serve as a sleep regulation hub: each SpsP neuron innervates all the PB glomeruli in the hemi-brain, receiving signals from and sending signals back to EB through EPG columnar neurons and PEcG columnar neurons, respectively, as well as by integrating dopaminergic signals.

## Results

### Calcium activity of protocerebral bridge (PB) increases with sleep deprivation

To address cellular mechanisms of sleep drive in a natural context, we took an unbiased snapshot of whole-brain calcium levels by pan-neuronally expressing CaMPARI. It is a photoconvertible calcium sensor, which irreversibly converts GFP to RFP in the presence of calcium and ultraviolet (UV) light^15,16^ (Fig. S1A). CaMPARI imaging has been validated and used in several studies to indicate the neuronal activity responses in flies to behaviors including olfaction^17^, bitter taste processing^18^ and hedonic hunger^19^. The RFP/GFP ratio after UV exposure is an indicator of relative calcium levels and therefore serves as a proxy for neuronal activity.

Flies exhibit rebound sleep after sleep deprivation^2,3^, supporting the notion that the sleep drive of an animal increases with wakefulness and dissipates with sleep. To identify the neurons that respond to high sleep drive, we carried out ex vivo imaging with pan-neuronally expressed CaMPARI at ZT1, i.e., one hour after lights on in a 12:12 LD cycle. This was done with two groups of flies: flies with high sleep drive that had been sleep deprived for 13 hours, i.e., for the whole night + 1 hr from ZT12 to ZT1 (SD group); and with flies harboring relatively low sleep drive which had slept undisturbed for the whole night and also assayed at the same time of day (CTRL group) (Fig. 1A). While whole brain CaMPARI imaging successfully labels neuropils throughout the entire brain, photoconversion within deep brain regions such as the EB and the FB might be limited by the penetration capability of the UV light and the laser for confocal imaging. Indeed, we found that the signal-to-noise ratios are weaker in neuropils deep in the brain compared to neuropils on the brain surface. We therefore normalized the CaMPARI signal of each SD neuropil to the same neuropil of the control group (see Methods for details). Among all the neuropils examined, the increase of CaMPARI signal after sleep deprivation in the PB was the most robust and reliable (Fig. 1B, Fig. S1C). The signal in the ellipsoid body (EB), a brain region that has been reported to encode sleep homeostasis^20^, also increased with sleep deprivation. There was little change in signal in the antenna lobe (AL) and the optic lobe (Fig. S1C).

**Figure 1.**
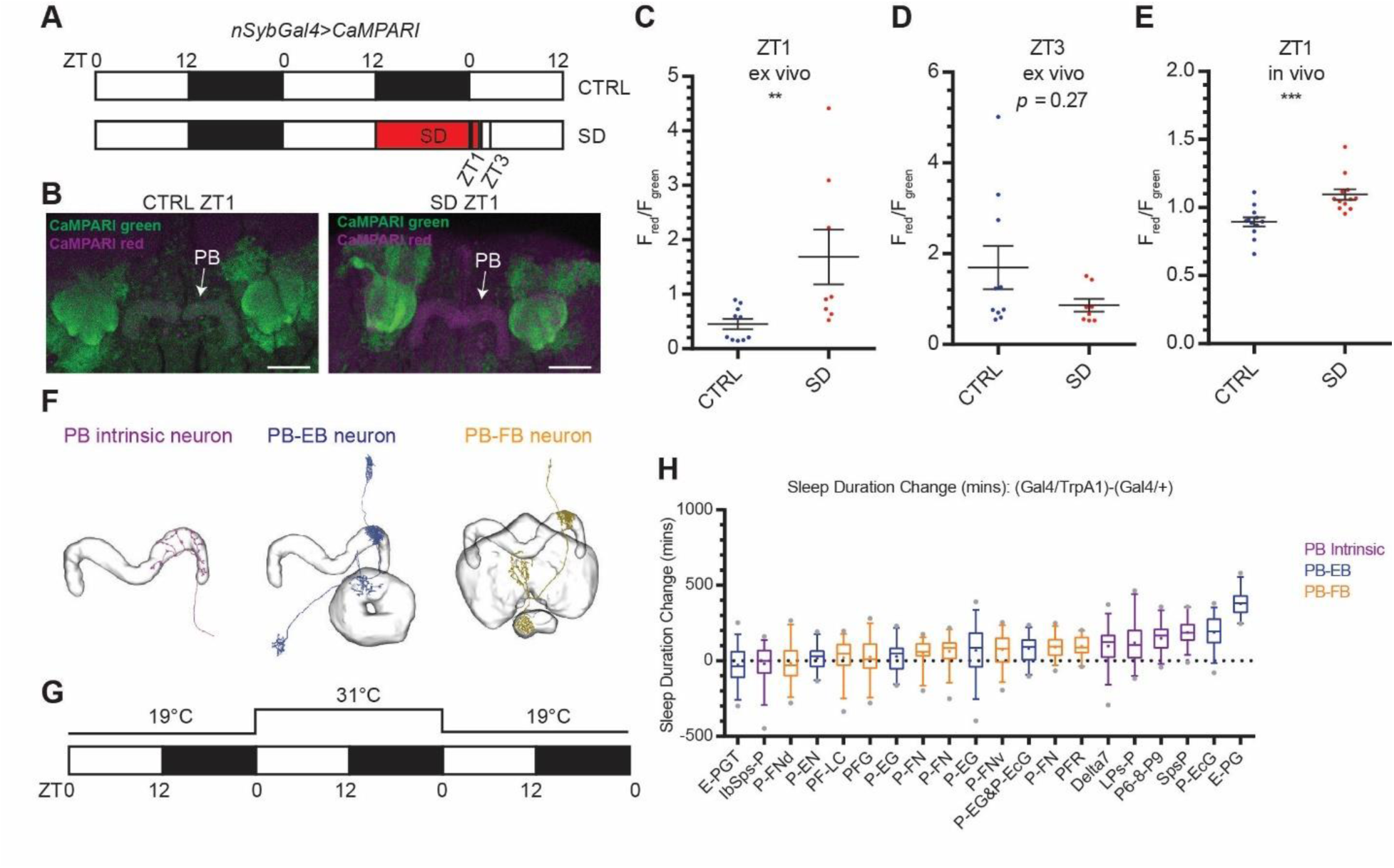
PB intrinsic and PB-EB neurons are involved in sleep regulation. **(A)** A schematic representation of the sleep deprivation paradigm for CaMPARI imaging. Male nSybGal4>CaMPARI flies were sleep deprived for 13 hours from ZT12 to ZT1 the next day for the sleep deprivation (SD) group. Flies in the SD group were sampled and imaged at ZT1 after 13hrs sleep deprivation and ZT3 after sleep deprivation followed by 2hrs of recovery sleep. Flies in the control (CTRL) groups were sampled and imaged at ZT1 and ZT3. Open bar: light phase, ZT0-12; solid bar: dark phase, ZT12-0; red bar: sleep deprivation. **(B)** Representative images of nSybGal4>CaMPARI imaging in the CTRL group (left) and the SD group (right) at ZT1. Green: CaMPARI Green; magenta: CaMPARI Red. Arrows: protocerebral bridge (PB). Scale bar: 30 µm. **(C-E)** Quantification of the F_red_/F_green_ ratio of nSybGal4>CaMPARI in the PB in the CTRL (blue) and the SD (red) group by ex vivo **(C-D)** and in vivo **(E)** CaMPARI imaging at ZT1 **(C, E)** and ZT3 **(D)**. Error bars: S.E.M.. ** P<0.001, ** P<0.01, Mann-Whitney test. **(F)** Representative projection patterns of three types of PB-projecting neurons. Images of representative neurons are from NeuPrint. **(G)** A schematic representation of the thermogenetic activation screen. The sleep of flies was monitored at 19°C during Day 1, followed by thermogenetic activation at 31°C during Day 2, and back at 19°C for recovery during Day 3. **(H)** Quantification of the sleep duration change when each of the 20 PB-projecting split-Gal4 drivers were thermogenetically activated by dTrpA1. Top three sleep-promoting drivers from right to left: E-PG (ss50574), P-EcG (ss02195), and SpsP (ss52267). Magenta boxes: PB-intrinsic neurons labeling drivers; Blue boxes: PB-EB neurons labeling drivers; Orange boxes: PB-FB neurons labeling drivers. More than 30 male flies were measured in each group. Whiskers represent 5 to 95 percentiles.

To further address the extent to which increased PB calcium after sleep deprivation is caused by increased sleep drive, we also carried out ex vivo CaMPARI imaging at ZT3 after 2 hours of recovery sleep (Fig. 1A). After 13 hours of sleep deprivation, flies in the SD group showed significantly higher PB calcium activity than the control group at ZT1 (Fig. 1C). And after additional 2 hours of recovery sleep at ZT3, there was no significant difference in the calcium activity of the PB between the SD group and the CTRL group (Fig. 1D). To further confirm the calcium activity of increase with sleep drive in the PB, we carried out in vivo CaMPARI imaging, in which freely moving flies were exposed to UV to capture ongoing calcium activity. In this assay as well, sleep deprived flies showed significantly higher PB calcium activity than the control group at ZT1 (Fig. 1E). These data indicate that PB calcium levels correlate with sleep drive.

### Thermogenetic activation screen of PB-expressing split drivers

The PB is a neuropil near the posterior surface of a fly brain, and it is part of the central complex. As mentioned above, other components of the central complex, most notably the FB^21^ and the EB^20,22,23^, also play important roles in sleep regulation. The sleep homeostasis process indicates that sleep drive increases with sleep deprivation and then leads to rebound sleep^24^. If increased PB calcium is due to sleep deprivation and reflects enhanced sleep drive, these neurons might promote sleep. But which are the relevant neurons?

There are about 3000 neurons in the central complex, of which about 660 neurons arborize in the PB^25^. They also connect to other central complex neurons to form highly-ordered neural networks. To understand which subsets of PB-expressing neurons function in sleep regulation and to avoid targeting non-specific neurons, we made use of the split-Gal4 library generated by Wolff et al.^26^. These lines have greatly improved specificity than the traditional driver lines because Gal4 is only active in the intersection of the p65AD and Gal4DBD patterns^27^. Based on their projection pattern/morphology, there are 3 categories of the PB-expressing neurons: 1) PB intrinsic neurons with most of their projections in the PB and no arborization in the EB or the FB; 2) PB-EB neurons, which innervate the PB and the EB; 3) PB-FB neurons, which innervate the PB and the FB (Fig. 1F).

To identify which sets of PB-expressing neurons function in sleep regulation, we carried out a thermogenetic activation screen with the PB-expressing split-Gal4 library (Fig. 1G); each driver labeled a sparse subset of PB-expressing neurons. When comparing the sleep change upon thermo-activation between activated Gal4>dTrpA1 flies and Gal4/+ control flies, no wake-promoting driver was found in our thermogenetic activation screen of PB-expressing drivers (Fig. 1H). This is consistent with the increased CaMPARI signal in the PB upon sleep deprivation. Sleep was significantly increased when several groups of PB-EB connecting neurons and PB intrinsic neurons were thermogenetically activated. In contrast, activation of PB-FB connecting neurons had only a minor or negligible effect (Fig. 1H), suggesting that the PB and its connection with the EB have a more important role in sleep regulation than the connection between the PB and the FB. The top 3 drivers with the most robust sleep-increasing phenotypes are ss52267, ss50574, and ss02195. Based on the brain regions they innervate, the relevant neurons labeled by these 3 drivers have been named by Wolff et al.^26^ as the SpsP neurons (innervating the superior posterior slopes (SPS) and the PB), the EPG neurons (innervating the EB, the PB, and the Gall), and the PEcG neurons (innervating the PB, the EB canal, and the Gall) respectively.

### Four SpsP neurons promote sleep

As the SpsP driver (ss52267) only labels four SpsP neurons in the central complex and weakly labels a few additional cells in the brain and the ventral nerve cord (VNC) (Fig. 2A, Fig. S2A), it is an attractive candidate for follow-up studies. Thermogenetic activation of the SpsP driver resulted in a daytime as well as a nighttime sleep increase (Fig. 2B-C). Notably, activation did not alter locomotor activity during movement compared to the TrpA1/+ control group (Fig. 2D), suggesting that the sleep increase is not due to defective locomotion. Thermogenetic activation of SpsP also led to more consolidated sleep; sleep bout duration increased (Fig. 2E), and sleep bout number decreased (Fig. 2F). In addition, optogenetic activation of the SpsP driver caused a nighttime sleep increase (Fig. S3A-B). Importantly, blocking vesicle endocytosis and thus synaptic transmission in the SpsP neurons by expressing shibire^ts28^ also caused sleep loss at night from ZT14 to ZT21 (Fig. 2G-H).

**Figure 2.**
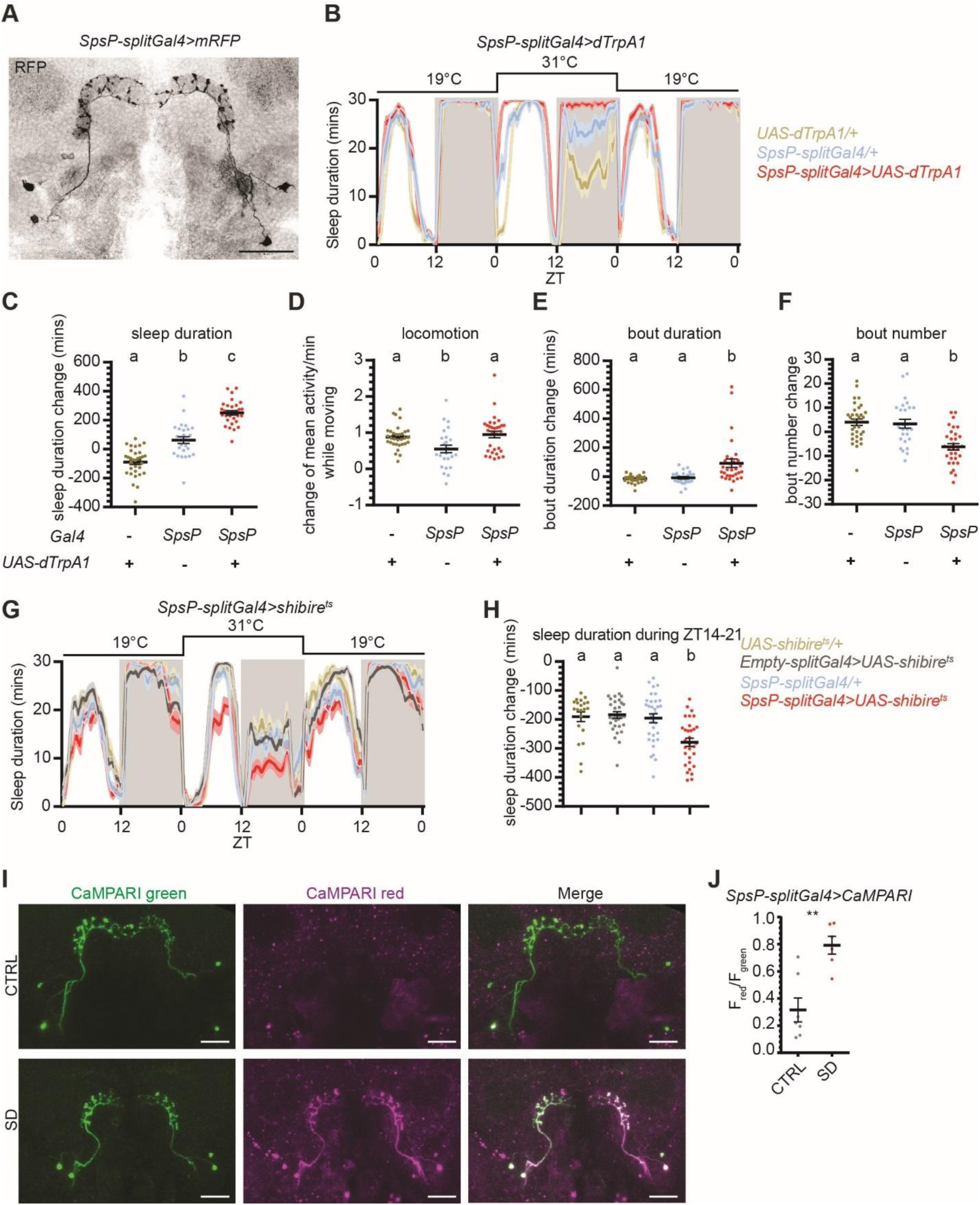
Four SpsP neurons promote sleep. **(A)** Representative image of the four SpsP cells labeled by SpsP-splitGal4>mRFP stained for RFP. Scale bar: 30 µm. **(B-F)** Sleep profiles **(B)** and quantification of the changes in sleep duration **(C)**, locomotion **(D)**, sleep bout duration **(E)**, sleep bout number **(F)** from ZT0-24 upon thermogenetic activation of the SpsP neurons. Sleep profiles are averaged in 30 minute bins. Changes were calculated by subtracting the value of baseline on Day 1 from that of activation on Day 2. Shaded area/Error bars: S.E.M.. Red: SpsP-splitGal4>UAS-dTrpA1 (n=31); Yellow: UAS-dTrpA1/+ (n=32); Blue: SpsP-splitGal4/+ (n=26). Letters represent statistically distinct groups; P < 0.01, Kruskal–Wallis test followed by a post hoc Dunn’s test. Male flies were used. **(G-H)** Sleep profiles **(G)** and quantification of the changes in sleep duration during ZT14-21 **(H)** upon thermogenetic inhibition of the SpsP neurons. Sleep profiles are averaged in 30 minute bins. Changes were calculated by subtracting the value of baseline from ZT14-21 on Day 1 from that of inhibition from ZT14-21 on Day 2. Shaded area/Error bars: S.E.M.. Red: SpsP-splitGal4>UAS-shibire^ts^ (n=31); Gray: Empty-splitGal4>UAS-shibire^ts^ (n=31); Yellow: UAS-shibire^ts^ /+ (n=21); Blue: SpsP-splitGal4/+ (n=32). Letters represent statistically distinct groups; P < 0.001, Kruskal–Wallis test followed by a post hoc Dunn’s test. Male flies were used. (**I**) Representative images of SpsP-splitGal4>CaMPARI flies in the CTRL (top) and the SleepDeprivation (SD) (bottom) group of flies. Green: CaMPARI Green; magenta: CaMPARI Red. Scale bar: 30 µm. (**J**) Quantification of the CaMPARI F_red_/F_green_ ratio in the cell bodies of the SpsP cells of SpsP-splitGal4>CaMPARI flies in the CTRL group (n=6) and the SD group (n=7). The average F_red_/F_green_ was calculated for each fly. Each dot represents an individual fly. Error bars: S.E.M.. **P<0.01, Mann-Whitney test. Male flies were used.

Based on the calcium increase in the PB with overnight sleep deprivation (Fig. 1C), we also assayed the SpsP neurons with CaMPARI imaging in flies after overnight sleep deprivation: calcium in the SpsP cell bodies increased after overnight sleep deprivation (Fig. 2I-J).

All of these results taken together indicate that the SpsP neurons are important for sleep and that their activity increases with sleep drive.

### SpsP neurons receive signals from and send signals to the PB

To explore the neural circuit basis of sleep regulation by SpsP neurons, we first took advantage of the recently published neuPrint analysis tool of fly brain electron microscopy (EM) reconstruction^29,30^. The EM reconstruction was carried out in a large portion of the central brain of the fly (or referred to as “hemibrain”). The 4 SpsP neurons are all annotated in neuPrint (ID: 881212361; 911893174; 941136727; 941469110), and the EM reconstruction shows that each SpsP neuron expands its projections across all nine PB glomeruli of each half hemibrain. Moreover, there are two SpsP neurons that project to the PB in the left side of the hemibrain, and two SpsP neurons that project to the PB in the right side of the hemibrain (Fig. 3A). However, the EM reconstruction is missing the SpsP arborizations within the SPS region, probably because it was not carried out on a full brain. Synaptic connectivity analysis based on the EM reconstruction only identifies sparse SpsP connections, most of which remain within the central complex: SpsP neurons receive inputs from upstream neurons in the PB, including Delta7 and SpsP itself, and they send outputs to downstream neurons in the PB, including PB intrinsic neurons such as Delta7 and SpsP, PB-FB neurons such as P-FNd, and PB-EB neurons such as PEN (Fig. 3B).

**Figure 3.**
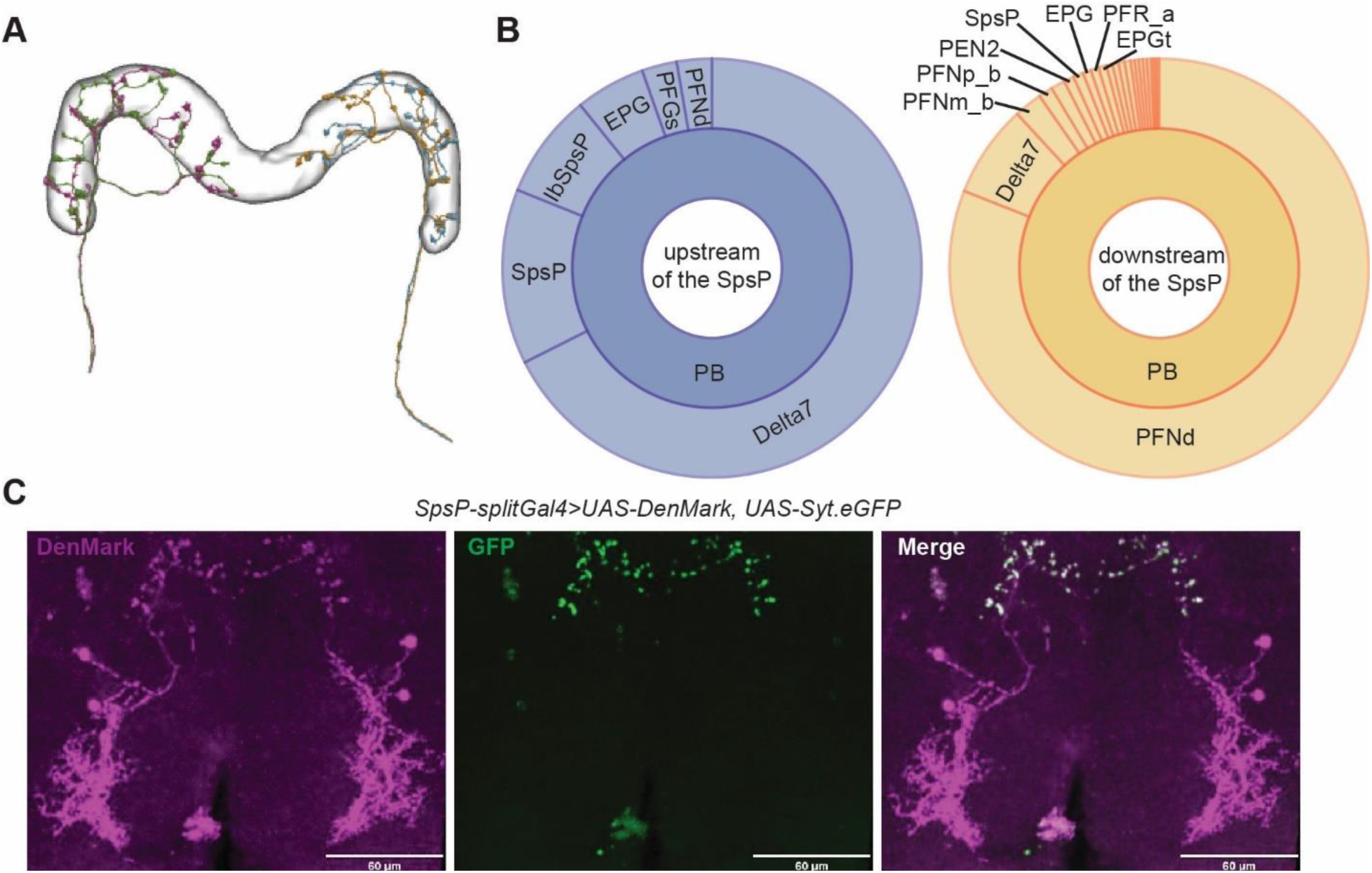
SpsP neurons receive signals from and send signals to the PB. **(A)** The projection patterns of the four SpsP neurons in the PB based on the electron microscopy (EM) reconstruction from NeuPrint. Each color represents an individual SpsP neuron. **(B)** Connectivity analysis of a representative SpsP neuron based on the EM reconstruction, adapted from NeuPrint. Left: Upstream brain regions and neurons of the representative SpsP neuron. Right: Downstream brain regions and neurons of the representative SpsP neuron. The size of each section represents the relative abundance of synaptic connections between the SpsP neuron and the specific type of neurons. **(C)** Representative images of the dendrites (left) and the axons (middle) of the SpsP cells labeled by SpsP-splitGal4>UAS-DenMark, UAS-Syt.eGFP stained for dsRed and GFP. Scale bar: 60 µm.

We then labeled the dendrites of SpsP neurons with DenMark^31^ and the axons with Syt.eGFP^32^. Similar to the results from the EM reconstruction, the immunostaining showed that SpsP neuron dendrites receive signals in the SPS and PB, and SpsP neuron axons send signals to the PB (Fig. 3C).

### SpsP neurons receive convergent activating signals from the EB via EPG neurons

Because both the EM reconstruction and immunostaining showed that most of the input and output signals of SpsP neurons lie within the PB, we postulated that most of the upstream and downstream neurons of SpsP neurons relevant to sleep regulation would be covered by our thermogenetic activation screen of PB-expressing split-Gal4 drivers. These neurons might include the EPG neurons, which caused the most robust sleep increase phenotype in the screen (Fig. 1H) and was indicated to lie upstream of the SpsP neurons by the connectivity analysis of the EM reconstruction (Fig. 3B).

Expression of mCD8-GFP^33^ in the EPG driver labels about 20 pairs of EPG neurons in the brain (Fig. 4A), with only 2 additional neurons weakly labeled in the VNC (Fig. S2B). Whereas the EPG driver labels all the glomeruli of the PB and the EB, EPG neurons are columnar neurons that tile together, so each individual EPG neuron is a columnar neuron that innervates one section of the EB, one of the 18 glomeruli of PB, and the gall (Fig. 4C). Immunostaining with DenMark and Syt.eGFP shows that the EPG dendrites are mostly in the EB and the axons in the PB (Fig. 4B), indicating that EPG neurons send signals from the EB to the PB. Consistent with this notion, connectomic analysis based on the EM reconstruction showed that EPG neurons lie upstream of SpsP neurons and form synapses with them. Thus, signals from different EPG neurons potentially converge onto each SpsP neuron via the PB neuropil.

**Figure 4.**
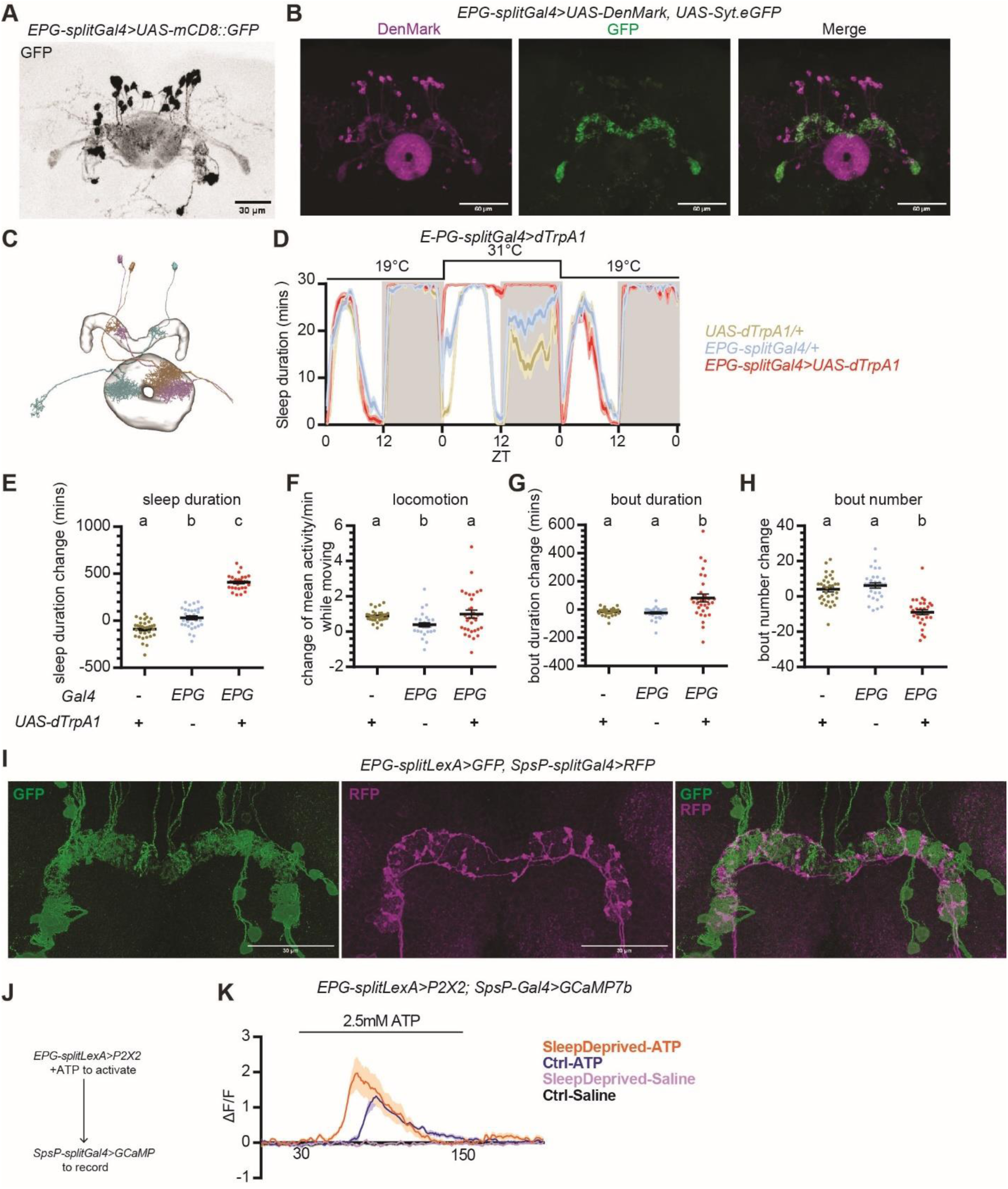
SpsP neurons are downstream of EPG neurons. **(A)** Representative image of the EPG neurons labeled by EPG-splitGal4>UAS-mCD8::GFP stained for GFP. Scale bar: 30 µm. **(B)** Representative images of the dendrites and the axons of the EPG neurons labeled by EPG-splitGal4>UAS-DenMark, UAS-Syt.eGFP stained for dsRed and GFP. Scale bar: 60 µm. **(C)** Projection patterns of three representative EPG neurons based on the EM reconstruction from NeuPrint. Each color represents an individual EPG neuron. **(D-H)** Sleep profiles **(D)** and quantification of the changes in sleep duration **(E)**, locomotion **(F)**, sleep bout duration **(G)**, sleep bout number **(H)** from ZT0-24 upon thermogenetic activation of EPG neurons. Sleep profiles are averaged in 30 minute bins. Changes were calculated by subtracting the value of baseline on Day 1 from that of activation on Day 2. Shaded area/Error bars: S.E.M.. Red: EPG-splitGal4>UAS-dTrpA1; Yellow: UAS-dTrpA1/+; Blue: EPG-splitGal4/+. Letters represent statistically distinct groups; P < 0.0001, Kruskal–Wallis test followed by a post hoc Dunn’s test. More than 30 male flies were used for each group. **(I)** Representative images of EPG-splitLexA>LexAop-mCD8::GFP, SpsP-splitGal4>UAS-IVS-mCD8::RFP stained for dsRed and GFP. Scale bar: 30 µm. **(J)** Schematic representation of the experimental design to test functional connectivity from the EPG neurons to the SpsP neurons. **(K)** Average GCaMP traces (ΔF/F) of SpsP cells in response to EPG activation. More than 5 male flies were used for each group. Error bars: S.E.M..

Thermogenetic activation of the EPG driver (ss50574) resulted in a dramatic increase in daytime and nighttime sleep duration (Fig. 4D-E) without significantly affecting locomotor activity (Fig. 4F). Similar to thermogenetic activation of the SpsP neurons, thermogenetic activation of the EPG neurons also resulted in less fragmented sleep, with the sleep bout duration significantly increased (Fig. 4G) and sleep bout number decreased (Fig. 4H). Optogenetic activation of the EPG neurons with CsChrimson expression similarly caused a consistent and robust sleep increase (Fig. S4A-B), indicating that the EPG neurons also promote sleep. However, blocking vesicle endocytosis with shibire^ts^ in EPG neurons had a puzzling effect by leading to a mild nighttime sleep increase rather than a decrease (Fig. S4C-D).

To further investigate the connection between the SpsP neurons and the EPG neurons, different components needed to be expressed in the two sets of neurons. To this end, we generated split-LexA lines to label these neurons; we verified that the new split-LexA lines had the identical expression patterns of the split-Gal4 lines by expressing LexAop-GFP and UAS-RFP simultaneously (Fig. S5).

Using the split-LexA line, GFP was expressed in the EPG neurons and RFP expressed in the SpsP neurons simultaneously. This revealed many adjacent projections of the EPG neurons and the SpsP neurons in the PB (Fig. 4I), indicating a likely synaptic connection between them.

We then assayed for a functional connection between the EPG neurons and the SpsP neurons by monitoring the levels of calcium and the second messenger cyclic AMP (cAMP) in the SpsP neurons upon activation of EPG neurons. To monitor cAMP levels, we generated a 20xUAS-EPACH187 line, which is more sensitive to cAMP than the original EPAC sensor^34–37^. GCaMP or EPACH187 was expressed in the SpsP neurons to record calcium or cAMP levels, and the ATP-gated cation channel P2X2 was expressed in the EPG neurons (Fig. 4J).

Activation of EPG neurons with ATP perfusion caused a significant increase in both the calcium level and the inverse FRET signal in the SpsP neurons (Fig. 4K, Fig. S4E), indicating a functional connection from the EPG neurons to the SpsP neurons. Moreover, the calcium response is even stronger in flies that have been sleep deprived for 13 hours overnight compared to the control group (Fig. 4K), suggesting that the connection is further strengthened by sleep deprivation.

### The connections from SpsP neurons to the PEcG neurons are dynamically modulated by sleep drive

Because the axons of the SpsP neurons are in the PB, we next asked which subset of PB-expressing neurons act downstream of the SpsP neurons to regulate sleep. Synaptic connectivity analysis based on the EM reconstruction indicated that one of the major cell types that lies downstream of the SpsP neurons is the P-FNd neurons (neurons that projects to the PB, the fan-shaped body, and the noduli), and SpsP neurons have been reported to act upstream of the P-FNd neurons in velocity tuning^38^. However, the sleep profile of P-FNd activation is comparable to the control groups (Fig. S6), suggesting that the SpsP neurons regulate sleep through a different circuit.

In our thermogenetic activation screen of PB-expressing split-Gal4 drivers, the PEcG driver (ss02195) showed the most robust sleep change among drivers that label neurons that receive signal inputs from the PB (Fig. 1H).

Expression of mCD8-GFP with the PEcG driver confirmed that it labels about 9 pairs of PEcG neurons (Fig. 5A) in the brain with nothing detectable in the VNC (Fig. S2C). PEcG neurons are canal P-EG neurons: a majority of their dendrites arborize in the PB, and their axons wrap the EB canal^26^ (Fig. 5B), sending signals from the PB to the EB. Like the EPG neurons, each individual PEcG neuron is a columnar neuron that innervates one section of the EB, one glomerulus of the PB, and the gall^26^. The PEcG neurons are not annotated by the EM reconstruction. Trans-synaptic mapping of the SpsP neurons using trans-Tango^39^ showed that the SpsP neurons send signals to neurons expressed in the EB and the PB (Fig. 5C), which is similar to the arborization pattern of the PEcG neurons. Moreover, the synaptic connections between the presynaptic sites of the SpsP neurons and the PEcG neurons were investigated by GFP Reconstitution Across Synaptic Partners (GRASP)^40^ imaging. Presynaptic nSyb-spGFP1-10 was expressed in the SpsP neurons and CD4-spGFP11 expressed in the PEcG neurons simultaneously. Immunostaining of reconstituted GFP showed synaptic connections between the SpsP neurons and the PEcG neurons (Fig. 5D), supporting the notion that the SpsP neurons send signals to the PEcG neurons.

**Figure 5.**
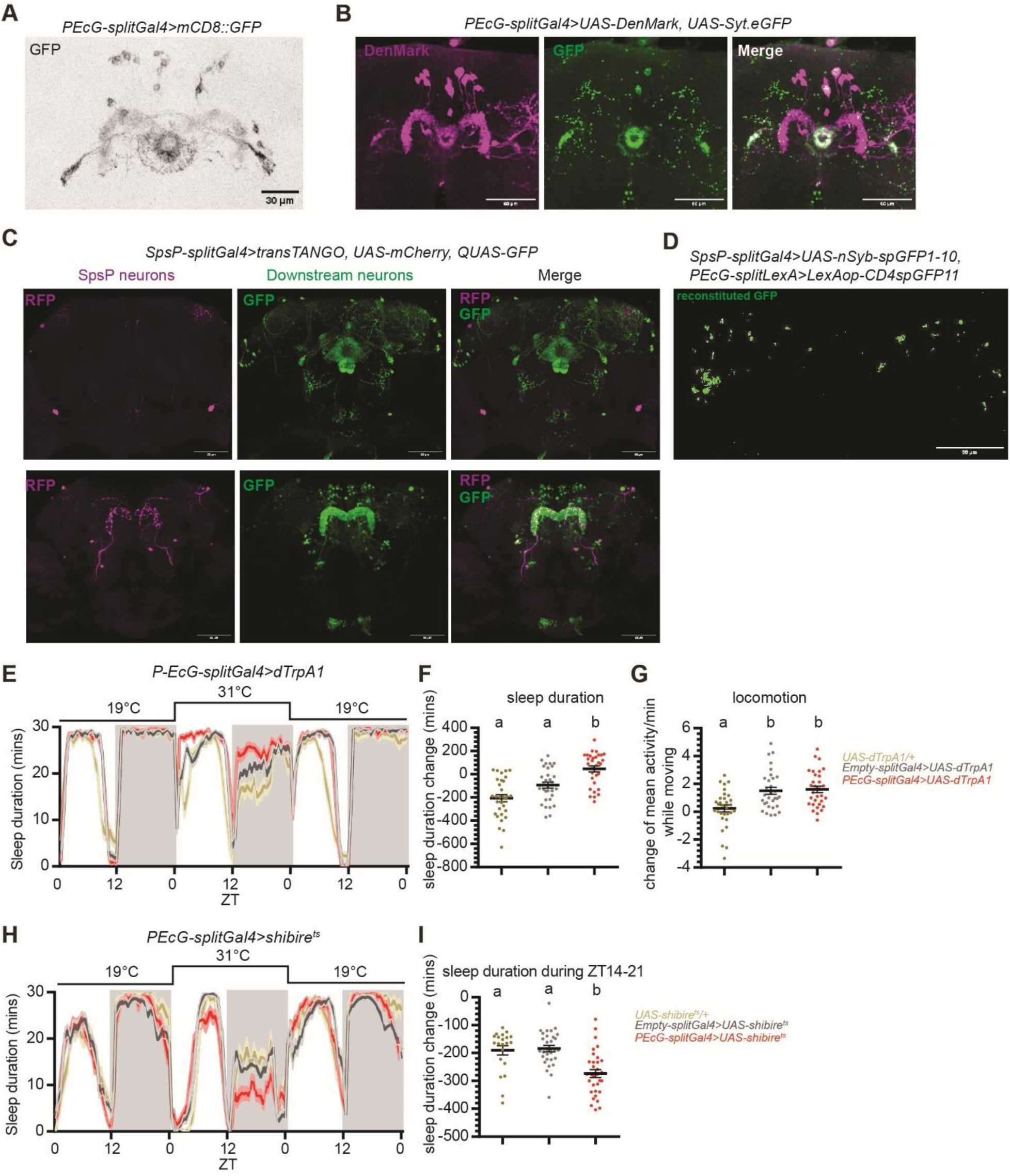
SpsP neurons are upstream of sleep-promoting PEcG neurons. **(A)** Representative images of the PEcG neurons labeled by PEcG-splitGal4>UAS-mCD8::GFP stained for GFP. Scale bar: 30 µm. **(B)** Representative images of the dendrites (magenta) and the axons (green) of the PEcG neurons labeled by EPG-splitGal4>UAS-DenMark, UAS-Syt.eGFP stained for dsRed and GFP. Scale bar: 60 µm. **(C)** Representative images of SpsP neurons (magenta) and their downstream neurons (green) labeled by SpsP-splitGal4>transTANGO, UAS-mCherry; QUAS-GFP stained for dsRed and GFP. Scale bar: 60 µm. **(D)** Representative GRASP image of the synaptic connections in the PB between SpsP and PEcG neurons labeled by SpsP-splitGal4>UAS-nSyb-spGFP1-10, PEcG-splitLexA>LexAop-CD4spGFP11 stained for reconstituted GFP. Scale bar: 60 µm. **(E-G)** Sleep profiles **(E)** and quantification of the changes in sleep duration **(F)** and locomotion **(G)** from ZT0-24 upon thermogenetic activation of the PEcG neurons. Sleep profiles are averaged in 30 minute bins. Changes were calculated by subtracting the value of baseline on Day 1 from that of activation on Day 2. Shaded area/Error bars: S.E.M.. Red: PEcG-splitGal4>UAS-dTrpA1; Gray: Empty-splitGal4>UAS-dTrpA1; Yellow: UAS-dTrpA1/+. Letters represent statistically distinct groups; P < 0.01, Kruskal–Wallis test followed by a post hoc Dunn’s test. More than 30 male flies were used for each group. **(H-I)** Sleep profiles **(H)** and quantification of the changes in sleep duration during ZT14-21 **(I)** upon thermogenetic inhibition of the PEcG neurons. Sleep profiles are averaged in 30 minute bins. Changes were calculated by subtracting the value of baseline from ZT14-21 on Day 1 from that of inhibition from ZT14-21 on Day 2. Shaded area/Error bars: S.E.M.. Red: PEcG-splitGal4>UAS-shibire^ts^ (n=32); Gray: Empty-splitGal4>UAS-shibire^ts^ (n=31); Yellow: UAS-shibire^ts^/+ (n=21). Letters represent statistically distinct groups; P < 0.001, Kruskal–Wallis test followed by a post hoc Dunn’s test. Male flies were used.

Like the SpsP neurons and the EPG neurons, thermogenetic activation of the PEcG neurons resulted in significant increases in both daytime and nighttime sleep duration (Fig. 5E-F), with no significant difference in locomotor activity (Fig. 5G). Optogenetic activation of the PEcG neurons also caused a sleep increase (Fig. S7), whereas blocking vesicle endocytosis in the PEcG neurons caused less nighttime sleep (Fig. 5H-I), indicating that the PEcG neurons contribute to sleep regulation.

We then examined the functional connection between the SpsP neurons and the PEcG neurons. Activation of the SpsP neurons by ATP perfusion led to a significant cAMP increase (Fig. S8) and calcium increase (Fig. 6A-B) in the PEcG neurons. Interestingly, the calcium response is further increased in sleep deprived flies (Fig. 6A-B), indicating that this functional connection between the SpsP neurons and the PEcG neurons is tunable.

**Figure 6.**
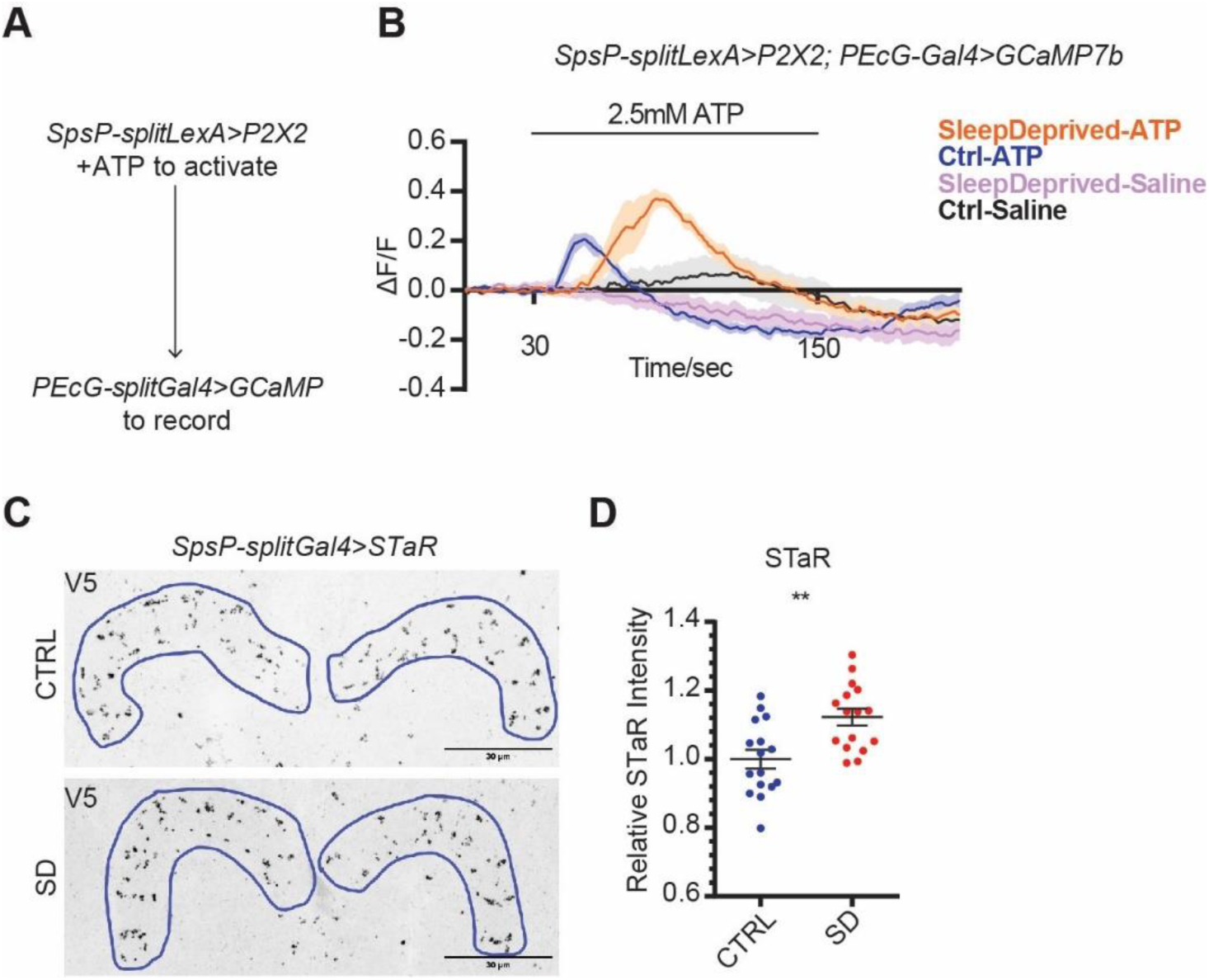
Sleep deprivation changes connectivity strengths between SpsP neurons and PEcG neurons. **(A)** Schematic representation of the experimental design to test functional connectivity from the SpsP neurons to the PEcG neurons. **(B)** Average GCaMP traces (ΔF/F) of PEcG cells in response to SpsP activation in the control group of flies (Ctrl, n=8) and flies after sleep deprivation (SleepDeprived, n=8). Male flies were used. Shaded area: S.E.M.. **(C-D)** Representative images **(C)** and quantification **(D)** of BRP protein abundance labeled by SpsP-splitGal4>UAS-Flp, brp(FRT.Stop)V5 stained for V5. Intensity was normalized to the average value of the control group. Examples of the regions of interest (ROIs) were drawn in blue. 8 male flies (16 hemi-PB ROIs) were used for each group. Scale bar: 30 µm. Error bars: S.E.M.. Mann-Whitney test, **P<0.01.

Are SpsP neurons altered upon sleep deprivation? Bruchpilot (BRP) is a key component of the presynaptic active zone, and the BRP abundance has been shown to correlate with vesicle release probability in *Drosophila*^41–44^. Using Synaptic Tagging with Recombination (STaR)^45^ to specifically tag the endogenous BRP protein with a V5 tag in the presynaptic sites of the SpsP neurons, we found that BRP is enriched in the SpsP neurons axons after sleep deprivation (Fig. 6C-D).

These results taken together indicate that the SpsP neurons act upstream of the PEcG neurons to promote sleep and that this connection is strengthened by sleep deprivation.

### SpsP neurons integrate dopaminergic signaling to regulate sleep

A previous single neuron labelling study with MultiColor FlpOut (MCFO) showed that individual SpsP neurons have abundant dendrites in the SPS^46^. This suggests that these neurons might integrate other sleep-relevant signals in addition to conveying signals between the EB and the PB. Because dopaminergic signaling is an important wake-promoting molecule in *Drosophila*^47–50^ and because several dopaminergic neuron clusters project to the SPS^51^, we asked whether the SpsP neurons might also integrate dopaminergic signals to regulate sleep.

To this end we first investigated whether the sleep-promoting SpsP neurons receive dopaminergic signaling. We made use of the DopR-Tango reporter^52^ that fuses the Tango system^53^ with UAS-DopR1 to report the sites of endogenous dopamine action. Expression of DopR-Tango driven by the SpsP driver confirmed that the SpsP cells receive dopaminergic signaling (Fig. 7A). Moreover, coexpression of GFP driven by SpsP-Gal4 and RFP driven by Dop2R-LexA shows that the two pairs of SpsP neurons colocalize with Dop2R-expressing neurons (Fig. 7B), whereas the other neurons labeled by the SpsP driver do not colocalize (Fig. S9A). This indicates that that the SpsP neurons express the inhibitory dopaminergic receptor Dop2R. Consistent with the sleep-promoting phenotype of activating the SpsP cells, knocking down Dop2R with RNAi (Fig. 7C-D) or mutating the gene with CRISPR-Cas9 (Fig. S9B-C) in these cells^54^ increased daytime sleep significantly. There was no increase in nighttime sleep possibly was not observed possible because of the ceiling effect. This supports an inhibitory role of Dop2R in the SpsP cells. Interestingly, rebound sleep was reduced after overnight sleep deprivation when Dop2R was downregulated (Fig. 7E-F, Fig. S9D-E).

**Figure 7.**
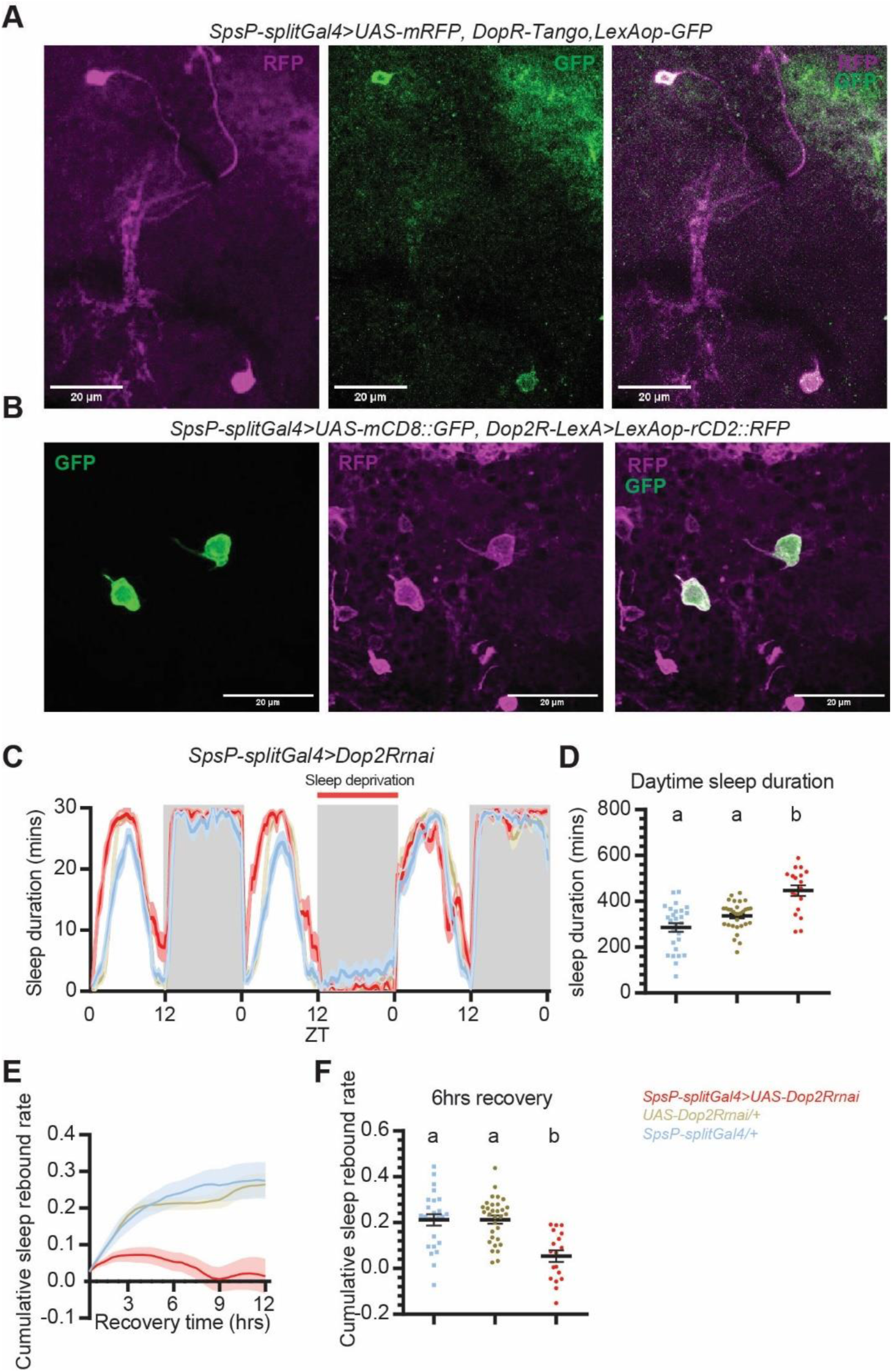
SpsP neurons integrate dopaminergic signaling to regulate sleep. **(A)** Representative images of the SpsP neurons (magenta) and the site of dopaminergic signaling (green) labeled by SpsP-splitGal4>UAS-mRFP, DopR-Tango, LexAop-GFP stained for dsRed and GFP. Scale bar: 20 µm. **(B)** Representative images of the SpsP neurons (green) and the Dop2R-expressing neurons (magenta) labeled by SpsP-splitGal4>UAS-mCD8::GFP, Dop2R-LexA>LexAop-rCD2::RFP stained for dsRed and GFP. Scale bar: 20 µm. **(C-F)** Sleep profiles **(C)**, quantification of the daytime sleep duration **(D)**, cumulative sleep rebound rate curve in 12 hours **(E)**, and the quantification of cumulative sleep rebound rate after 6 hours of recovery sleep **(F)** in the experimental group and the control groups of Dop2R knockdown in the SpsP neurons. **(C)** and **(E)** are average sleep duration in 30 minute bins. Cumulative sleep rebound rate was calculated by first dividing the sleep rebound by the sleep loss for each fly, and then cumulated over the bins. Shaded area/Error bars: S.E.M.. Red: SpsP-splitGal4>UAS-Dop2Rrnai (n=18); Yellow: UAS-Dop2Rrnai/+ (n=31); Blue: SpsP-splitGal4/+ (n=27). Letters represent statistically distinct groups; P < 0.01, Kruskal–Wallis test followed by a post hoc Dunn’s test. Virgin female flies were used.

We interpret these results to indicate that dopamine signaling acts like a gate in the SpsP cells to regulate sleep. During regular daytime, the SpsP cells are inhibited by wake-promoting dopaminergic neurons through Dop2R to help keep flies awake, whereas this Dop2R effect is suppressed after sleep deprivation. This helps activate SpsP cells to drive rebound sleep. Dop2R downregulating causes the SpsP cells to mimic a “rebound sleep” state and prevents them from mounting a greater sleep rebound response. The results taken together indicate that the SpsP neurons integrate dopaminergic signaling through Dop2R to further refine sleep regulation.

## Discussion

We have shown in this study that the activity of the PB increases with sleep deprivation and that the four key SpsP neurons that innervate this neuropile are sleep-promoting. To carry out this function, these SpsP neurons also communicate with the EB and do so by sending signals through the PEcG neurons and by receiving signals from the EPG neurons (Fig. 8). Because the SpsP neurons also receive important wake-promoting inputs via dopamine signaling, we suggest that the SpsP neurons are a new central player in fly brain sleep regulation.

**Figure 8.**
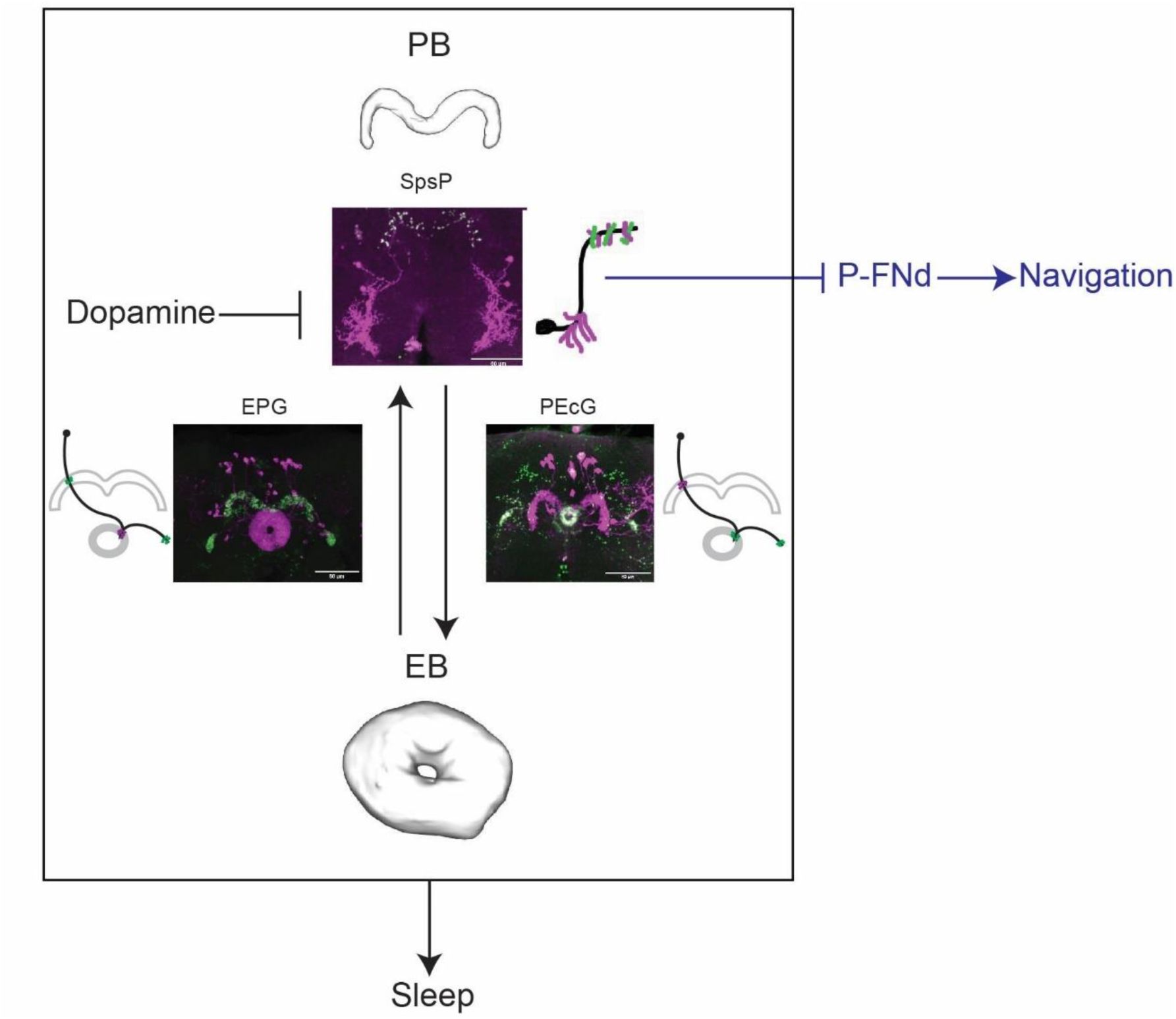
Summary of the SpsP circuit in sleep regulation and navigation. Projecting though the PB glomeruli, the SpsP neurons integrate signals from the EB through the EPG neurons and inhibitory dopaminergic signaling probably from the SPS, and send signals to the EB through the PEcG neurons to regulate sleep. The SpsP neurons also regulate navigation functions such as velocity tuning by inhibiting the P-FNd neurons^38,74^. Dendrites that receive signals are in magenta, axons that send signals are in green. Confocal images are DenMark and Syt.eGFP expressed in the split driver. Cartoons represent the projection patterns, dendrites, and axons, of individual neurons.

### An unbiased whole brain calcium snapshot captures active brain regions in sleep-deprived flies

Through analysis of CaMPARI signal in multiple brain regions, the PB, along with the well-known sleep regulating brain regions the EB and the MB, showed a robustly increased calcium level in sleep deprived flies. There was in contrast only a minimal effect in the vision processing optic lobe and the olfactory processing antenna lobe (Fig. S1C). This revealed the PB as a new player in sleep regulation. The calcium activity increase in the PB with sleep drive was observed in both ex vivo and in vivo CaMPARI imaging (Fig. 1C-E).

Whole brain CaMPARI imaging has two major advantages relative to other calcium indicators and should be able to identify networks that regulate specific internal states or behaviors different from sleep drive. 1) It is irreversible, allowing the capture of calcium activity snapshots that compare groups of flies of different internal states without any external transient stimuli such as drug application or neuronal manipulation. 2) It is non-invasive, enabling the in vivo imaging of freely moving flies, in contrast to the restricted mobilization used for traditional in vivo calcium imaging.

### A sleep promoting circuit between PB and EB inside the central complex

PB, EB, and FB are the three major components of the central complex. Several groups of central complex neurons, including dorsal FB neurons, helicon cells, and EB ring neurons, have been implicated in sleep regulation^20,21,50,55–59^. However, the role of PB neurons in sleep regulation has not been as extensively explored. Activation of PB-expressing drivers has only recently been shown to regulate sleep^60^, but this study was limited by the broad expression patterns of the Gal4 drivers, within as well as outside the PB.

Here, we show that as few as four SpsP neurons are important for normal sleep regulation. This probably reflects their role in integrating signals from, and conveying signals to, larger groups of neurons. Indeed, recent characterization by light microscopy^25,26,46,61–64^ and EM reconstruction^65^ showed that the fly central complex is a highly organized structure: there are columnar cells for parallel information processing and orthogonal tangential cells to collect inputs from and send outputs to these and other columnar cells. Each EPG and PEcG columnar neuron arborizes onto one section of the EB and one glomerulus of the PB. In contrast, the axons and dendrites of the tangential SpsP neurons cover all 9 pairs of PB glomeruli; these four neurons therefore serve as an integrating hub, by collecting inputs from the ∼40 EPG columnar neurons and sending output to the ∼18 PEcG columnar neurons.

We present several lines of evidence supporting the relevance to sleep of these collecting and sending functions of the SpsP neurons. 1) There was a significant sleep increase when the SpsP, the EPG, or the PEcG neurons were activated (Fig. 2B-C, Fig. 4D-E, Fig. 5E-F). Although inhibition phenotypes were weaker than activation, sleep levels were consistently decreased when synaptic transmission was blocked in the SpsP or the PEcG neurons (Fig. 2G, Fig. 5H). Notably, previous studies have shown sleep increases induced by activating other central complex sleep regulators like the dFB or EB-R5 neurons, but to our knowledge inhibiting these neurons has not been reported to cause sleep effects. Unexpectedly, blocking synaptic transmission in the EPG neurons caused increased sleep (Fig. S4C-D), probably because they are functionally more heterogenous than the less numerous SpsP and PEcG neurons. Consistent with this interpretation, the variation of locomotion and sleep bout duration is bigger in EPG-activated flies than the control groups (Fig. 4F-G). 2) The dendrites of the SpsP neurons (Fig. 3C) and the axons of the EPG neurons (Fig. 4B) are juxtaposed, as are the axons of the SpsP neurons (Fig. 3C) and the dendrites of the PEcG neurons (Fig, 5B). 3) TransTango imaging showed that the downstream neuron targets of the SpsP neurons innervate the EB and PB (Fig. 5C). 4) GRASP imaging showed synaptic connections between the SpsP neurons presynaptic sites and PEcG neurons within the PB (Fig. 5D). 5) EPG neuron activation increased SpsP calcium (Fig. 4K) and cAMP levels (Fig. S4E), and SpsP activation increased PEcG calcium (Fig. 6B) and cAMP levels (Fig. S8).

The circuit of EPG-SpsP-PEcG identified here suggests that information flow from PB-to-EB as well as its reverse, EB-to-PB, functions in sleep regulation. This substantially expands the understanding of how the central complex regulates sleep. The EB consists of tangential neurons known as ring neurons, which arborize in concentric rings and include the EB-R1, R2, R3, R4, and R5 neurons. There are also columnar neurons such as the EPG and PEcG neurons. EB-R5 neurons (originally termed as R2 neurons) are proposed to encode sleep drive^20,59^ and integrate signals from the circadian system^66–68^; they are also proposed to act upstream of dFB neurons^20^, which then inhibit helicon cells and also send signals back to the EB-R5 neurons^55^. This suggests a recurrent circuit for sleep homeostasis regulation of which the dFB neurons are the output effector arm. Other subtypes of EB ring neurons have also been identified in the regulation of sleep amount and sleep structure^22,69,70^. For example, R3m neurons are sleep-promoting and R3d neurons are wake-promoting.

The EB-R5 sleep regulating circuit is probably connected to the SpsP centric circuit we describe here: The EB-R5 neurons act upstream of the EPG neurons to regulate sleep and sleep homeostasis: sleep deprivation promotes EB-R5 synchronization, increases EPG neuron activity, and strengthens the connection between the EB-R5 and the EPG neurons^20,23,59^. Given the role of EPG-to-SpsP connections identified here, a reasonable scheme is that the EPG columnar neurons are conveying signals from the EB-R5 tangential neurons to the SpsP tangential neurons. The PEcG neurons, on the other hand, act downstream of the SpsP tangential neurons and send signals back to the EB. The PEcG neurons project broadly to the EB concentric rings, especially to the center or to the “canal” of the ring, and therefore have the potential to convey signals from the SpsP neurons to several groups of sleep-regulating EB ring neurons within the EB.

### SpsP neurons receive dopaminergic signaling to regulate sleep

In addition to communicating signals with the EB, the SpsP neurons respond to other inputs. DopR-Tango imaging shows that dopaminergic signaling impacts these neurons (Fig. 7A). Dop2R-LexA labels the four SpsP neurons but not the other cells of the SpsP split Gal4 driver (Fig. 7B, Fig. S9A). Moreover, Dop2R downregulation within SpsP neurons increased daily sleep and decreased recovery sleep after sleep deprivation (Fig. 7C-F, Fig. S9B-E), indicating that SpsP neurons probably receive inhibitory signals from dopaminergic neurons to reduce sleep. Might the inhibitory dopaminergic signal come from the PB, perhaps from the one pair of T1/LPs-P dopaminergic neurons that innervates PB^25,26,46,71^? Probably not, as we found that both thermogenetic and optogenetic activation of the LPs-P neurons (labeled by ss52578) caused a sleep increase rather than a decrease, and optogenetic inhibition of the LPs-P neurons showed no sleep effect (Fig. S9F-H). This indicates that if a connection exists between the LPs-P neurons and SpsP neurons, it is excitatory, i.e., the opposite of what is indicated by the Dop2R KO phenotype. The SpsP neurons more likely receive dopaminergic input from within the SPS, where their arbors coincide with those from PAL, PPL2ab, and PPL2c dopaminergic neurons^51^.

### Integration of sleep and navigation

The 4 SpsP neurons share two uncommon features with the EPG neurons: 1) Their activity increases with sleep deprivation, and they are sleep-promoting; 2) They are also involved in navigation. Indeed, EPG neurons are compass neurons that represent heading direction^72,73^, and SpsP neurons function in velocity tuning^38^. This function of SpsP neurons and that of sleep-promotion occur through distinct downstream circuits: the neurons send signals to the FB through the P-FNd neurons to regulate velocity tuning^38,74^ and send signals to the EB through PEcG neurons to regulate sleep. How does the central complex and more specifically its 4 SpsP neurons coordinate these two very different functions? One possibility is that they are subject to plasticity gating as a function of sleep and wake^75^. Indeed, our data indicate that enhanced sleep drive caused multiple changes in the SpsP circuit: After sleep deprivation, the calcium activity of all four SpsP neurons increased (Fig. 2I-J), BRP protein is more abundant in the SpsP axons (Fig. 6C-D), and the functional connections of EPG-to SpsP and SpsP-to-PEcG are stronger (Fig. 4J-K, Fig. 6A-B). Moreover, the connections of EBR5-to-EPG have been reported to increase with sleep deprivation^23^. We suggest that these changes integrate sleep and navigation in opposite directions: 1) They turn down navigation when an animal is about to sleep. The increase of activity and BRP abundance in the EBR5-EPG-SpsP circuit caused by sleep deprivation is accompanied by strong inhibition of the P-FNd neurons; this impairs velocity tuning and visual signal processing. 2) Given the known role of the EPG neurons and SpsP neurons in navigation, the opposite gate might be enhanced by extensive navigation. For example, when the animal encounters a new environment, it might strengthen the connection between the SpsP and P-FNd neurons at the expense of the SpsP-PEcG connection and sleep. Perhaps the compact nature of the small fly brain necessitates such switches. Verifying these gates and then understanding the underlying switching mechanisms are fascinating topics for future study.

## Materials and Methods

### Fly Stocks and Rearing

All flies were raised at 25°C on standard cornmeal food with a 12hr:12hr light: dark cycle. The genotypes of fly strains used are listed in Table S1.

### Generation of Fly Lines

To generate the split-LexA lines, pBPZpGAL4DBDUw (Addgene #26233) was firstly digested with KpnI (NEB #R3142S) and HindIII (NEB #R0104S), and then Gal4DBDZp was replaced with LexADBDZp amplified from UAS-LexADBD^76^ by Gibson assembly (NEB #E2611S) to generate a pBPLexADBDZp construct. The corresponding enhancer of each split driver was respectively amplified from the genomic DNA of wildtype flies, and ligated into AatII (NEB #R0117S) and NaeI (NEB #0190S) digested pBPLexADBDZp construct. Sequencing-verified enhancer-LexADBDZp plasmids were injected into attP1 site on the second chromosome by Rainbow Transgenic Flies Inc (Camarillo, CA, USA). Positive transformants were screened by eye color and confirmed by PCR. Enhancer-LexADBDZp flies were then recombined with corresponding p65ADZp lines that were inserted at attP40 site on the second chromosome to make a stable split-LexA line.

To generate the UAS-EPACH187 line, mCD8::GFP was cut out from pJFRC7-20XUAS-IVS-mCD8::GFP (Addgene #26220)^77^ by NotI and XbaI, and replaced with EPAC-S-H187 from Epac-S-H187 (Addgene #170348)^34^. Sequencing-verified UAS-EPACH187 plasmids were injected into attP1 site on the second chromosome by Rainbow Transgenic Flies Inc (Camarillo, CA, USA). Positive transformants were screened by eye color and confirmed by PCR.

### Sleep monitoring and analysis with DAM system

Flies of 5-10 days old were collected and loaded into glass behavioral tubes containing food of 5% sucrose and 2% agar. The tubes were then put onto the Drosophila Activity Monitors (DAM; TriKinetics Inc., Waltham, MA) system in an incubator of 25°C unless noted otherwise. Locomotion and sleep were analyzed by the 2020 version of SCAMP MATLAB Program.

### Thermogenetic neuronal manipulation

Flies were collected, loaded into the DAM system, and entrained at 19°C with a 12hr:12hr light: dark cycle. Baseline sleep was measured at 19°C for at least one day, and then transferred to 31°C for thermogenetic activation or inhibition. The temperature was then reduced back to 19°C for recovery measurements. Changes were calculated by subtracting the value of the baseline from the manipulation, and then compared between groups.

For the thermogenetic activation mini-screen of PB-expressing neurons, the sleep duration change was calculated and plotted by first subtracting the sleep durations on Day 2 (activation) by the sleep durations on Day 1 (baseline) for each fly, and then the average sleep duration change of the Gal4/+ control group was subtracted from each fly in the corresponding Gal4/dTrpA1 group.

### Sleep deprivation

Flies were collected and loaded into the DAM system, and then the DAM boards were loaded onto a vortexer mounting plate (TriKinetics Inc., Waltham, MA) mounted to a VWR shaker. Flies were vortexed for 2 seconds every ∼20 seconds (uniformly distributed randomized) during the sleep deprivation period^78^. Cumulative rebound rate of each individual fly was calculated as 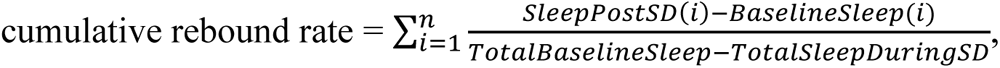 where i is the bin being analyzed, and sleep was analyzed in 30 minute bins, with 48 bins in each day. TotalBaselineSleep was calculated as the total sleep duration of the corresponding ZT time to sleep duration on the baseline day. For example, when sleep is deprived from ZT12 to ZT0, sleep duration from ZT12 to ZT0 on the day before was counted as TotalBaselineSleep.

### Optogenetic neuronal manipulation

Optogenetic experiments were carried out in the video-recording FlyBox^79^. Fly food consisting of 5% sucrose, 2% agar, and 400µM all trans-retinal (ATR) was dispensed into 96 well plates. 40mM ATR stock was made by dissolving the ATR powder (Sigma) in ethanol and stored in the dark at -20°C. Flies were transferred to the 96 well plates four days before the optogenetic stimulation. Each fly was put in an individual well. Flies were entrained with 12hr:12hr light:dark at 5 lux. Red light (LEDSupply, 624nm to 634nm, 0.1 mW mm^-2^) pulsing at 5Hz was used for CsChrimson experiments. Green light (LEDSupply, 520nm to 540nm, 0.1 mW mm^-2^) was used for GtAcr1 experiments. The activity of the flies was captured every 10 seconds by WebCamImageSave (NirSoft). Locomotion were then analyzed by the PySolo software^80^, trimmed into DAM readable format, and sleep was analyzed by the 2020 version of Sleep and Circadian Analysis MATLAB Program (SCAMP)^81^.

### Immunohistochemistry and confocal imaging

Flies at 5-10 days old were dissected in 1xPBS, the brains and ventral nerve cord were then fixed in 4% paraformaldehyde in PBS for 55 minutes. Tissues were then washed 3 x 10 minutes in 0.5% Triton X-100 in PBS (PBT), blocked in PBT with 10% normal goat serum(NGS) at 4°C overnight, incubated with primary antibodies in PBT with 10% NGS at 4°C overnight. Tissues were then washed 3 x 15 minutes in PBT, incubated with corresponding secondary antibodies in PBT with 10% NGS at 4°C overnight. Next, tissues were washed 3 x 15 minutes in the PBT and 15 minutes in the PBS, mounted in VectaShield mounting medium, and imaged using Leica Stellaris 8 confocal microscope with a white light laser. Images were processed and analyzed with ImageJ.

For GFP and RFP immunostaining, chicken anti-GFP (Abcam #ab13970; 1:1000) and rabbit anti-dsRed (Takara Bio #632393; 1:300) were used.

For GRASP experiments, the reconstituted GFP was immunostained with rabbit anti-recombinant GFP (Invitrogen #G10362; 1:100) and AlexaFluor488 anti-rabbit (Invitrogen #A-11008; 1:400).

For STaR experiments, the V5 tag was immunostained with mouse anti V5-tag conjugated to DyLight 550 (BioRad #MCA1360D550GA; 1:400).

### CaMPARI imaging

For ex vivo CaMPARI imaging, brains were first dissected in adult hemolymph like saline (AHL), UV light (LEDSupply, 400nm to 410nm) was then illuminated onto freshly dissected brains rinsed in AHL. The brains were then imaged using Leica Stellaris 8 confocal microscope. The UV light was pulsing in a 500ms ON, 200 ms OFF cycle (Fig. S1B, top), this cycle was on for 60 seconds and then off for 30 seconds with a duration of 150 seconds in total to avoid overheating (Fig. S1B, bottom) for a total duration of 150 seconds. The paradigm was verified by high K+ buffer application or dTrpA1 expression and activation.

For in vivo CaMPARI imaging, flies were put into a sylgard plate covered by a piece of qPCR film, holes were poked in the film to allow the flies to breathe. The UV light was pulsing in the same pattern as ex vivo CaMPARI imaging but with a duration of 900 seconds. Flies were then dissected, brains fixated in 4% PFA for 2hours, washed three times with wash buffer for 15minutes each time, mounted onto a glass slide, and imaged.

To compare the fluorescence signals between samples of different conditions, same settings of laser and detectors were used during each experiment. Sleep of each fly was first analyzed and picked before imaging, flies that had normal sleep were picked for the CTRL group, and flies that were sleep deprived were picked for the SD group. The signals were quantified by ImageJ and Fred/Fgreen was calculated using the method in Edward et al.^17^. Briefly, background was subtracted from each channel and then the Fred/Fgreen ratio was calculated.

### GCaMP Imaging

Male flies of 5-10 days old was used. Brains were dissected in ice-cold AHL, and then mounted onto small pieces of poly-L-Lysine-coated coverslips (Neuvitro #GG-12-PLL) on SYLGARD 184-coated Siskiyou perfusion chambers (Automate Scientific). ValveBank II perfusion system (Automate Scientific) was used to set up the perfusion flow. Briefly, solutions of AHL, ATP, and 85mM K+ buffer were put in individual channels. The flow rate was set to 1 drop per second. The brain was bathed in AHL before the focal plane was set. Each brain was put through three sets of flow consequentially: 1) AHL-AHL-AHL for a saline control, switching between two channels of AHL; 2) AHL-2.5mM ATP-AHL, switching between a channel of AHL and a channel of 2.5mM ATP solution flow for stimulation, followed by AHL flow for recovery and wash;3) AHL-85mM K+ buffer as a positive control. Only brains that were responding to 85mM K+ buffer were selected for analysis of the saline control and the response to 2.5mM ATP stimulation.

UAS-GCaMP7b and UAS-mRFP were recombined and imaged at the same time, mRFP serving as a endogenous control. Images were captured 1 frame per second on Leica Stellaris 8 confocal microscope, analyzed with custom MATLAB program modified from Adel et al.^82^. The change of mRFP over time was subtracted from GCaMP for correction. ΔF/F0 of the GCaMP signal was then calculated as (F-F0)/F0, where F is the signal at the given frame, and F0 is the baseline signal which is defined as the average signal of the first 10 frames.

### cAMP imaging

Male flies of 5-10 days old were used for cAMP imaging using the EPAC FRET sensor. The perfusion protocol was similar with that of GCaMP imaging, but 20uM Forskolin (cAMP agonist) was used as positive control instead of 85mM K+ buffer. Only brains that were responding to 20uM Forskolin were selected for analysis of the saline control and the response to 2.5mM ATP stimulation.

For analysis, background-subtracted CFP and YFP signals were measured for every frame, and the inverse FRET (iFRET) signals were calculated with methods modified from Shafer et al.^35^ to reflect the cAMP level: F= CFP/ (YFP-(CFP*0.27)), where 0.27 is the measured CFP spillover into the YFP channel in our scope settings. ΔF/F of the iFRET signal was then calculated as (F-F0)/F0, where F is the signal at the given frame, and F0 is the baseline signal which is defined as the average signal of the first 10 frames.

### Statistical Analysis

Statistical tests were performed using the Statistics Kingdom website.

## Acknowledgements

We thank all the Rosbash lab members, especially Shlesha Richhariya, E. Nicholas Petersen, and Katherine Abruzzi, for the discussion and help for this work. We thank Fang Guo, Enxing Zhou, Renbo Mao, Leslie Griffith, and Paul Garrity for helpful discussions and comments on the manuscript. We thank Katie Lyons for the assistance in maintaining the fly stocks. We thank Mohamed Adel for the help in building up the setup for functional imaging. We thank Chi-Hon Lee for the UAS-LexADBD construct. We thank Heather Dionne and Gerry Rubin (Janelia Research campus) for the UAS-shibire^ts^ fly line, and Yi Rao for the Dop2R-LexA fly line. Flies from Bloomington *Drosophila* Stock Center were used for this study. This study is supported by the Howard Hughes Medical Institute (HHMI).

## Author contributions

Conceptualization: XD, MR. Methodology: XD, JL, MR. Investigation: XD, JL, DM, MR. Supervision: MR. Writing—original draft: XD, MR. Writing—review & editing: XD, JL, DM, MR.

## Supplementary Materials

### Supplementary Figures

**Fig. S1.**
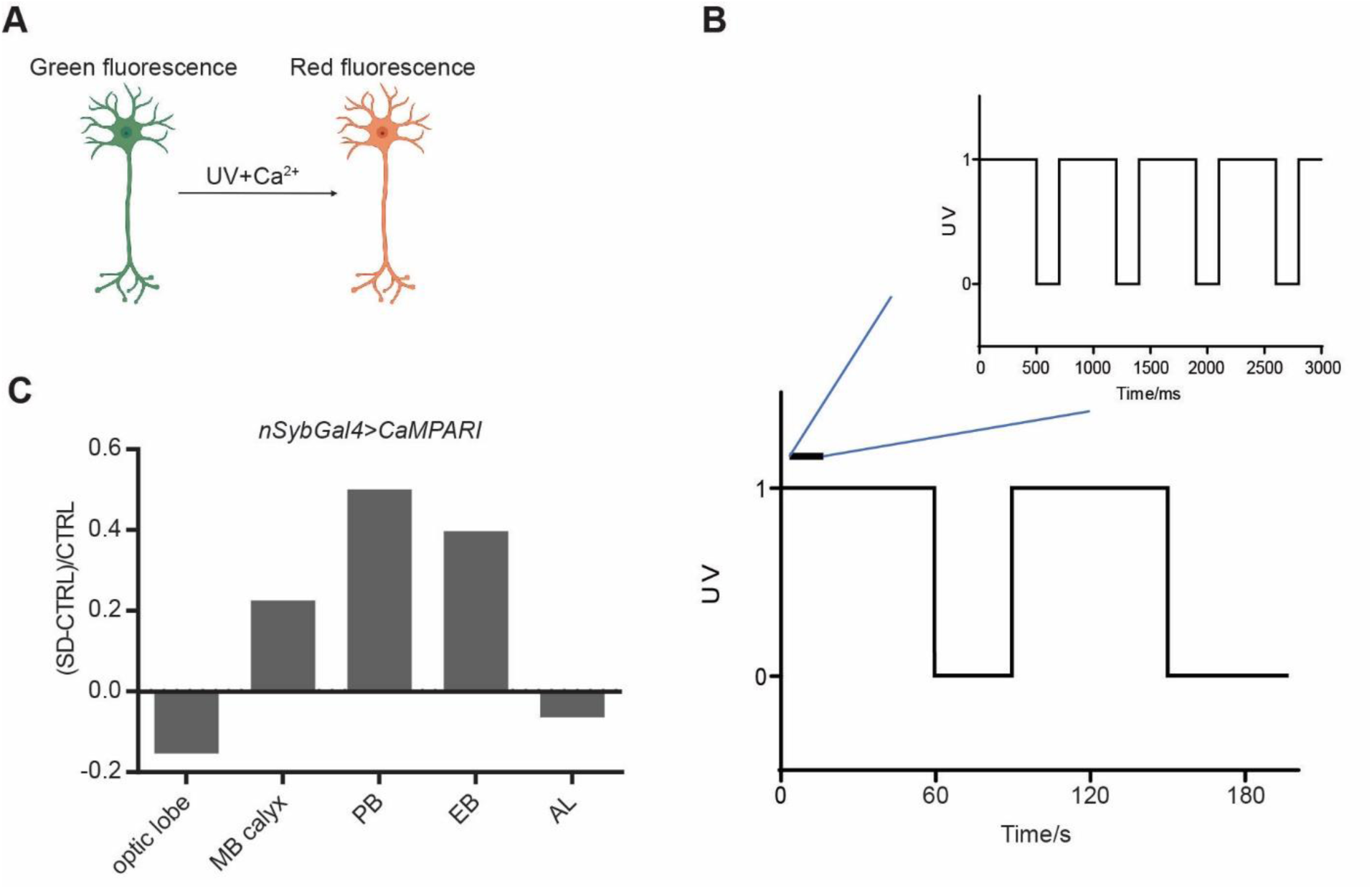
Calcium activity changes in different brain regions after sleep deprivation. **(A)** A schematic representation of CaMPARI imaging. CaMPARI converts from green to red fluorescence in the presence of UV and high concentration of calcium. **(B)** Paradigm of UV pulsing in CaMPARI imaging. UV is pulsing with 500ms ON and 200ms OFF for 60 seconds followed by 30 seconds of break until a total duration of 150 seconds is reached. **(C)** Normalized change of CaMPARI F_red_/F_green_ ratio between the SD group after 13hrs sleep deprivation and the CTRL group at ZT1 in different brain regions.

**Fig. S2.**
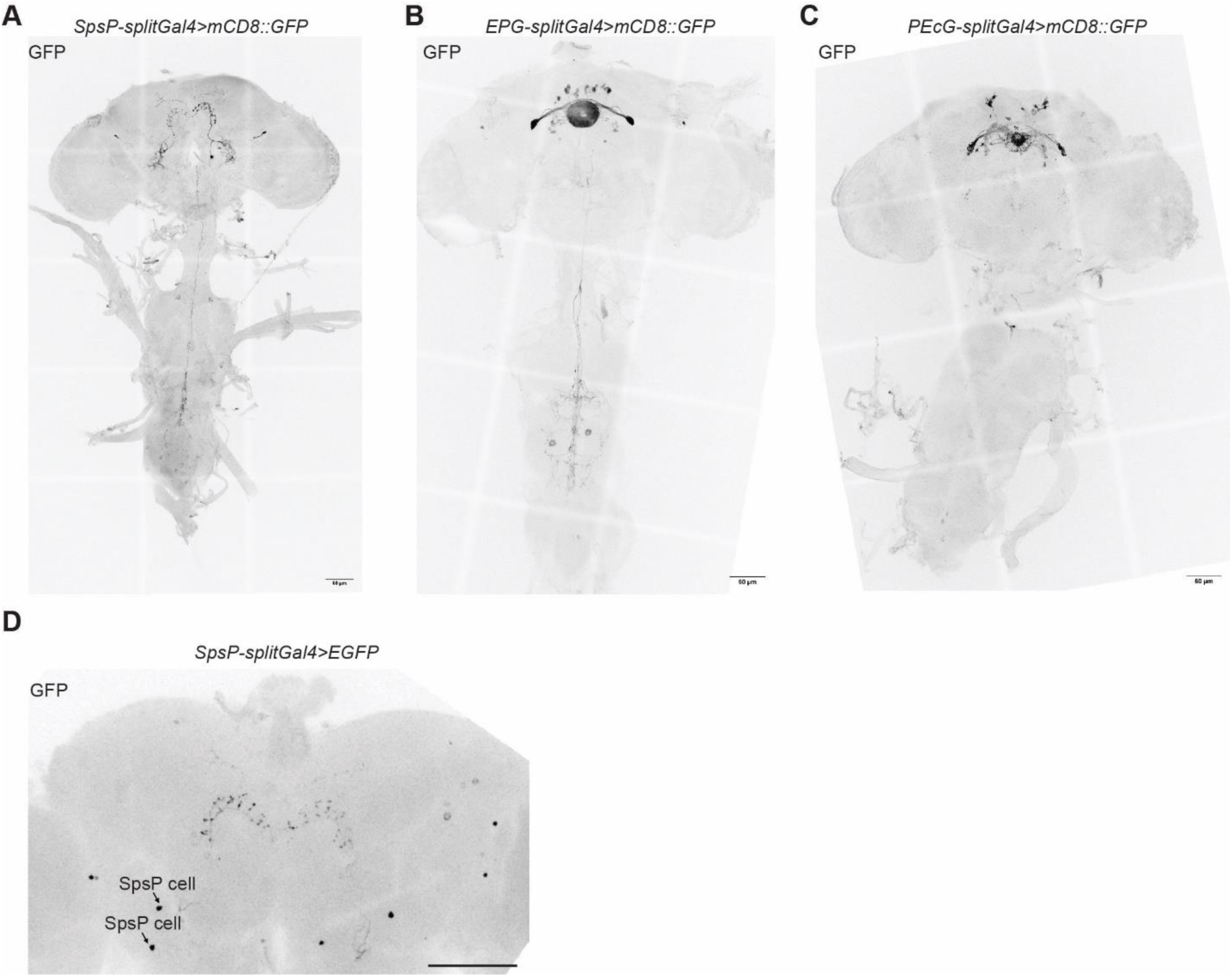
Candidate split drivers show sparse and specific expression patterns. **(A-C)** Representative images of SpsP-splitGal4(ss52267)>mCD8::GFP **(A)**, EPG-splitGal4(ss50574)>mCD8::GFP **(B)**, and PEcG-splitGal4(ss02195)>mCD8::GFP (**C)** stained for GFP in the brain and the VNC. Scale bar: 60 µm. **(D)** Representative image of SpsP-splitGal4>EGFP stained for GFP in the brain. Arrows: Cell bodies of the SpsP cells in the left half-brain. Scale bar: 60 µm.

**Fig. S3.**
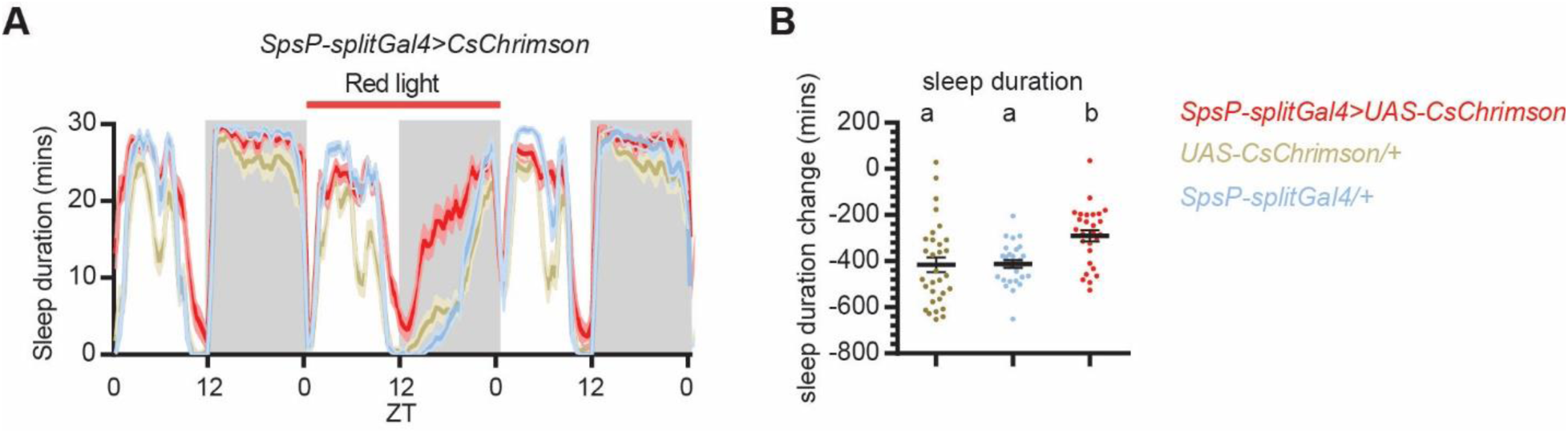
Optogenetic activation of SpsP neurons promote sleep. **(A-B)** Sleep profiles **(A)** and quantification of the changes of sleep duration **(B)** from ZT0-24 upon optogenetic activation of the SpsP neurons. Sleep profiles are averaged in 30 minute bins. Changes were calculated by subtracting the sleep duration of Day 1 ZT0-24 from that of Day 2. Shaded area/Error bars: S.E.M.. Red: SpsP-splitGal4>>UAS-CsChrimson (n=28); Yellow: UAS-CsChrimson/+ (n=31); Blue: SpsP-splitGal4/+ (n=29). Letters represent statistically distinct groups; P < 0.01, Kruskal–Wallis test followed by a post hoc Dunn’s test. Male flies were used. Male flies were used.

**Fig. S4.**
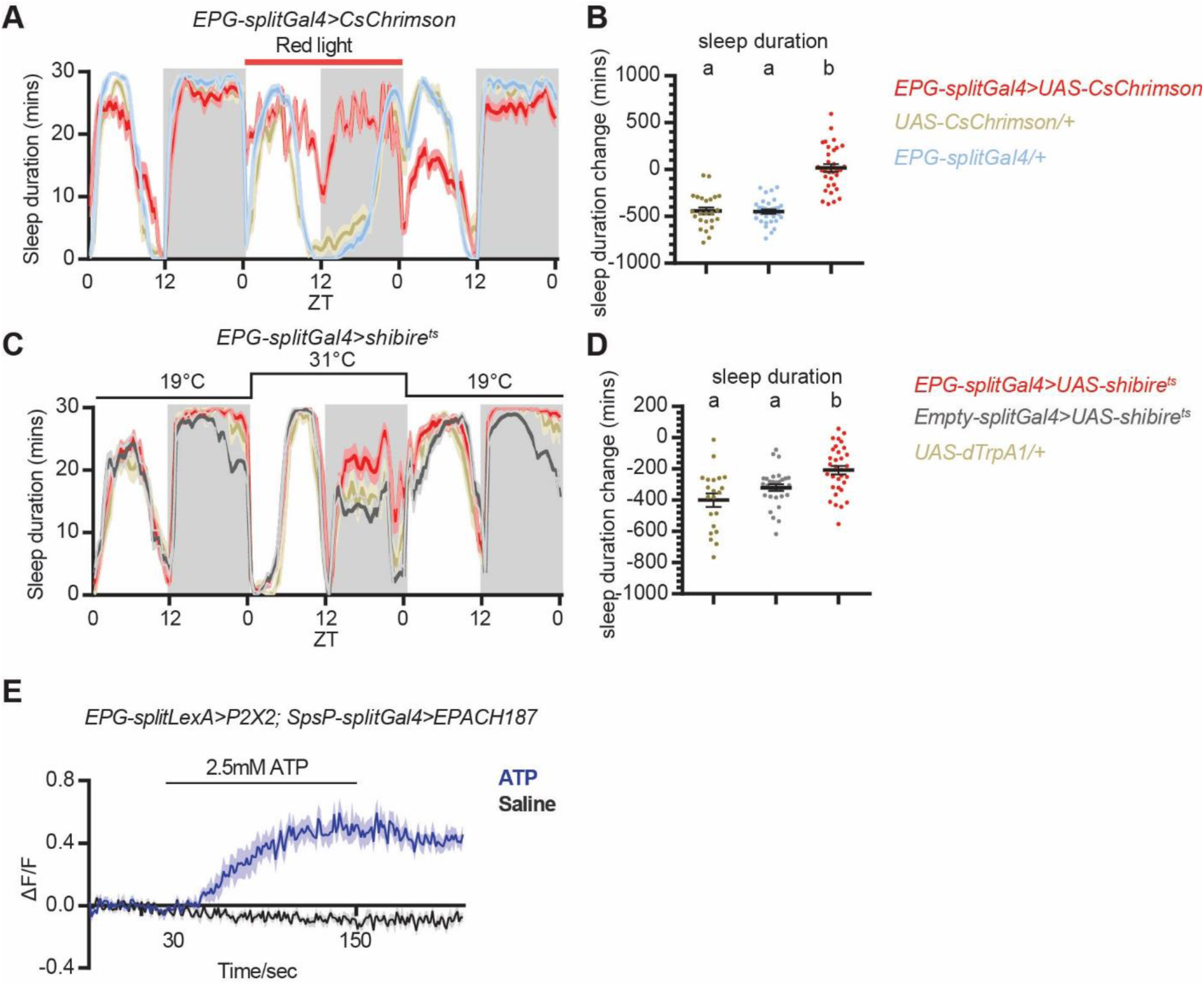
Activation of EPG neurons causes increased sleep and increased ATP level in SpsP neurons, whereas inhibition of E-PG neurons causes mildly increased nighttime sleep. Sleep profiles **(A, C)** and quantification of the changes in sleep duration **(B, D)** from ZT0-24 upon optogenetic activation **(A)** or neurotransmitter release blocking **(C-D)** of the EPG neurons. Sleep profiles are averaged in 30 minute bins. Changes were calculated by subtracting the sleep duration of Day 1 ZT0-24 from that of Day 2. Shaded area/Error bars: S.E.M.. Red: EPG-splitGal4/UAS; Yellow: UAS/+; Blue: EPG-splitGal4/+; Gray: Empty-splitGal4/UAS. More than 20 male flies were used for each group. Letters represent statistically distinct groups; P < 0.05, Kruskal–Wallis test followed by a post hoc Dunn’s test. **(E)** Average EPAC traces (ΔF/F: inverse FRET signal) of SpsP cells in response to EPG activation. More than 7 male flies were used for each group. Error bars: S.E.M..

**Fig. S5.**
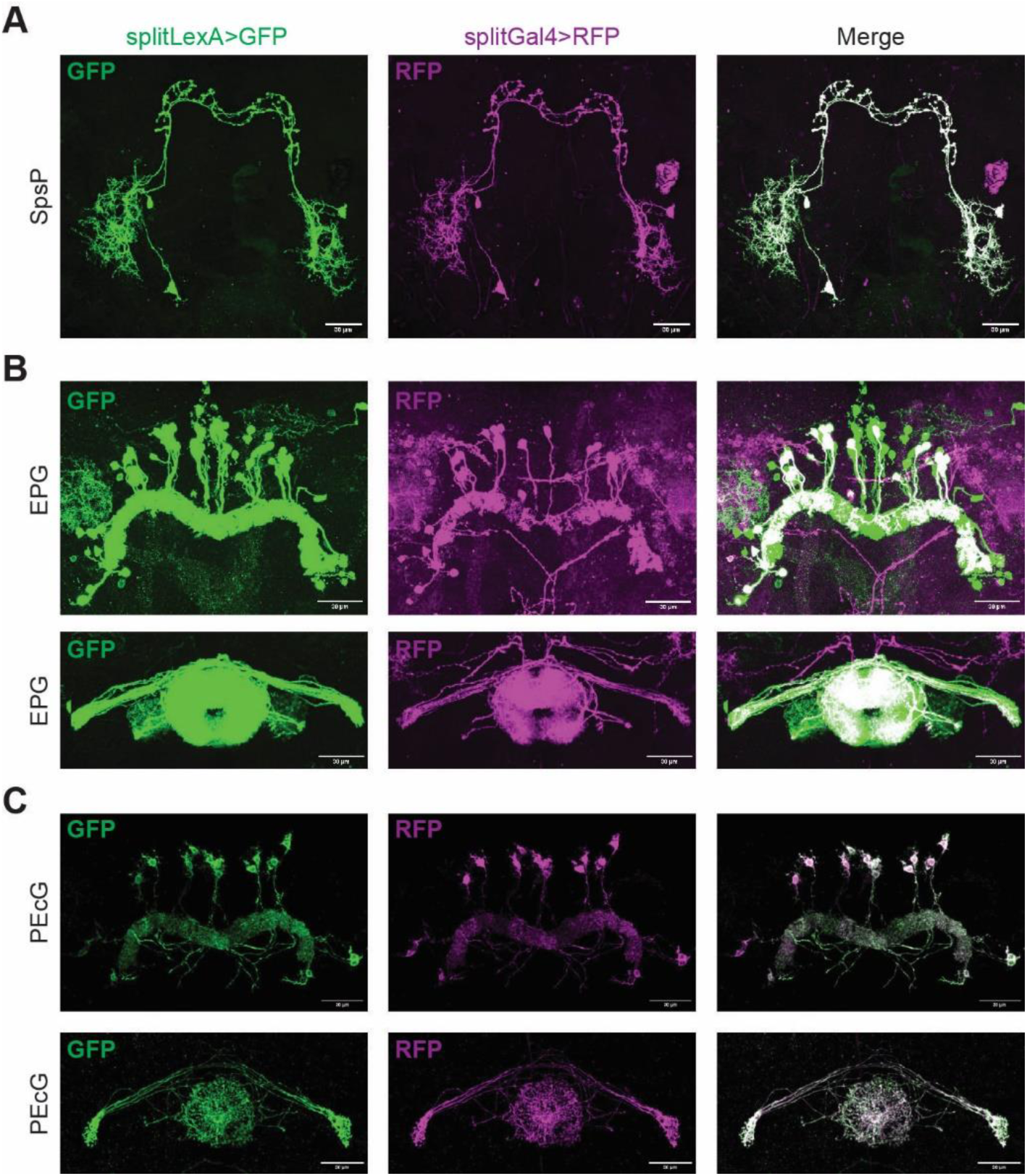
The expression patterns of split-LexAs recapitulate those of corresponding split-Gal4s. Representative images of neurons labeled by split-LexA drivers (left) and split-Gal4 drivers of SpsP **(A)**, EPG **(B)**, and PEcG **(C)** stained for GFP and dsRed. Scale bar: 30 µm.

**Fig. S6.**
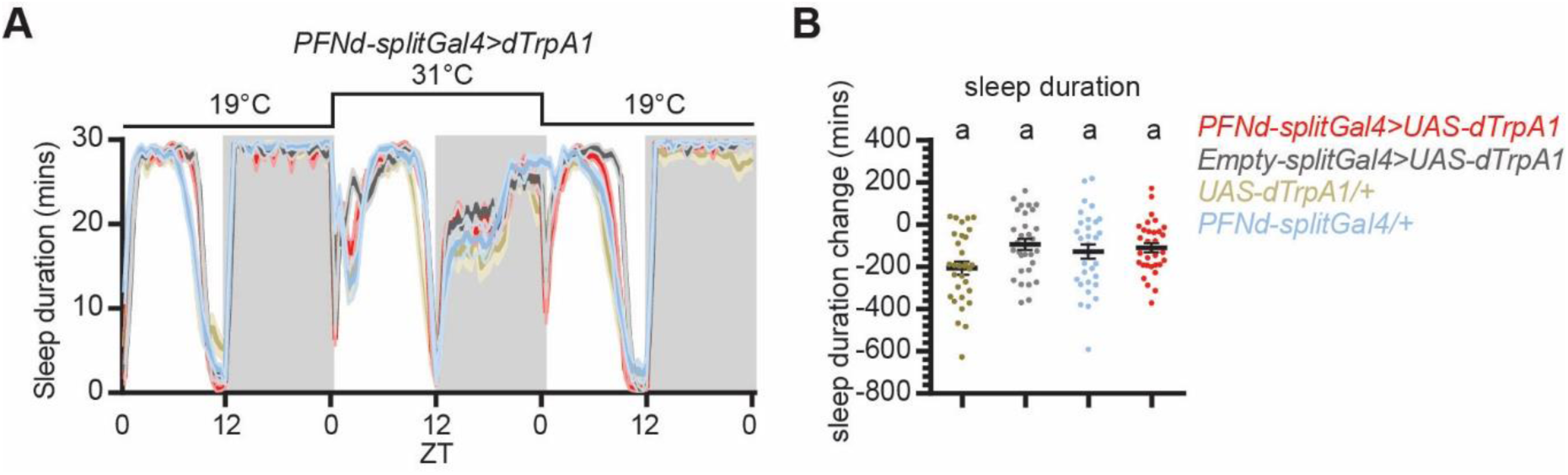
Activation of the PFNd neurons has no effect in sleep duration. Sleep profiles **(A)** and quantification of the changes in sleep duration **(B)** from ZT0-24 upon thermogenetic activation of the PFNd neurons. Heatshock was from Day2 ZT0-24. Sleep profiles are averaged in 30 minute bins. Changes were calculated by subtracting the sleep duration of Day 1 ZT0-24 from that of Day 2. Shaded area/Error bars: S.E.M.. Red: PFNd-splitGal4>UAS-dTrpA1 (n=32); Gray: Empty-splitGal4/UAS-dTrpA1 (n=31); Yellow: UAS-dTrpA1/+ (n=31); Blue: PFNd-splitGal4/+ (n=31). Same letters represent statistically indistinct groups; P > 0.05, Kruskal– Wallis test followed by a post hoc Dunn’s test. Male flies were used.

**Fig. S7.**
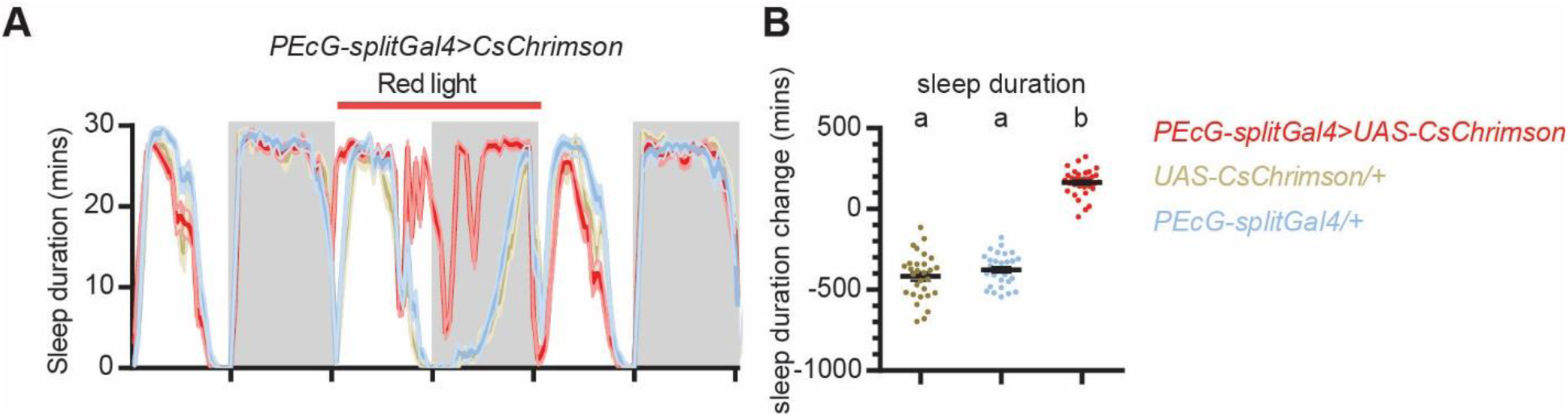
Activation of the PEcG neurons causes a sleep increase. Sleep profiles **(A)** and quantification of the changes in sleep duration **(B)** of ZT0-24 upon optogenetic activation of the P-EcG neurons. Sleep profiles are averaged in 30 minute bins. Shaded area/Error bars: S.E.M.. Changes were calculated by subtracting the sleep duration of Day 1 ZT0-24 from that of Day 2. Red: P-EcG-splitGal4>UAS-CsChrimson (n=32); Yellow: UAS-CsChrimson/+ (n=29); Blue: P-EcG-splitGal4/+ (n=30). Letters represent statistically distinct groups; P < 0.0001, Kruskal– Wallis test followed by a post hoc Dunn’s test.

**Fig. S8.**
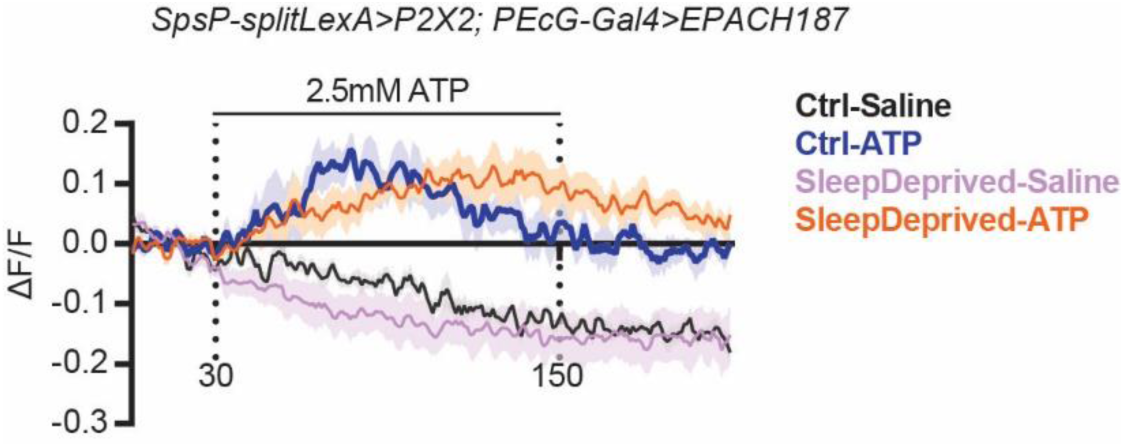
Activation of the SpsP neurons causes cAMP level increase in the P-EcG neurons. Averaged EPAC traces (ΔF/F, F: inverse FRET signal) of PEcG cells in response to ATP-induced SpsP activation. n=5 male flies for each group. Error bars: S.E.M..

**Fig. S9.**
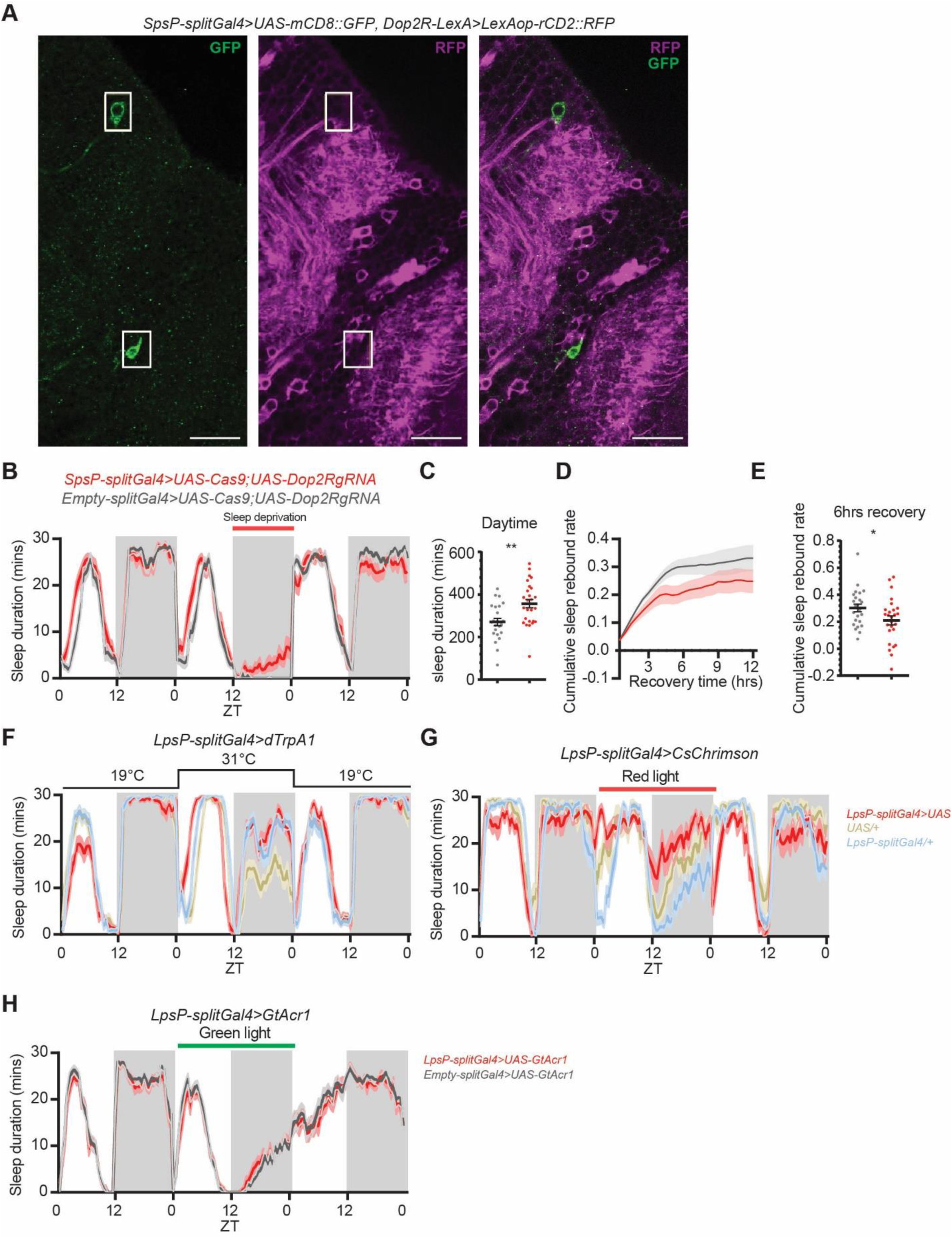
SpsP neurons integrate dopaminergic signaling to regulate sleep. **(A)** Representative images of the additional neurons labeled by the SpsP driver (green) and the Dop2R-expressing neurons (magenta) labeled by SpsP-splitGal4>UAS-mCD8::GFP, Dop2R-LexA>LexAop-rCD2::RFP stained for dsRed and GFP. Scale bar: 20 µm. **(B-E)** Sleep profiles **(B)**, quantification of the daytime sleep duration from ZT0-12 **(C)**, cumulative sleep rebound rate curve in 12 hours **(D)**, and the quantification of cumulative sleep rebound rate after 6 hours of recovery sleep **(E)** in the experimental group and the control groups of SpsP specific Dop2R mutated flies. **(B)** and **(D)** are in 30 minute bins. Shaded area/Error bars: S.E.M.. Red: SpsP-splitGal4>UAS-Cas9; UAS-Dop2RgRNA (n=26); Gray: Empty-splitGal4>UAS-Cas9; UAS-Dop2RgRNA (n=24). *P<0.05, **P<0.01, Mann-Whitney test. Virgin female flies were used. **(F-H)** Sleep profiles of thermogenetic activation **(F)**, optogenetic activation **(G)**, and optogenetic inhibition **(H)** of the dopaminergic LpsP neurons in 30 minute bins. Red: LpsP-splitGal4/UAS; Yellow: UAS/+; Blue: LpsP-splitGal4/+; Gray: Empty-splitGal4/UAS. More than 20 male flies were used for each group.

