## Supplemental figures 1-9 for "Four SpsP neurons are an integrating sleep regulation hub in *Drosophila*"

### Supplementary Figures

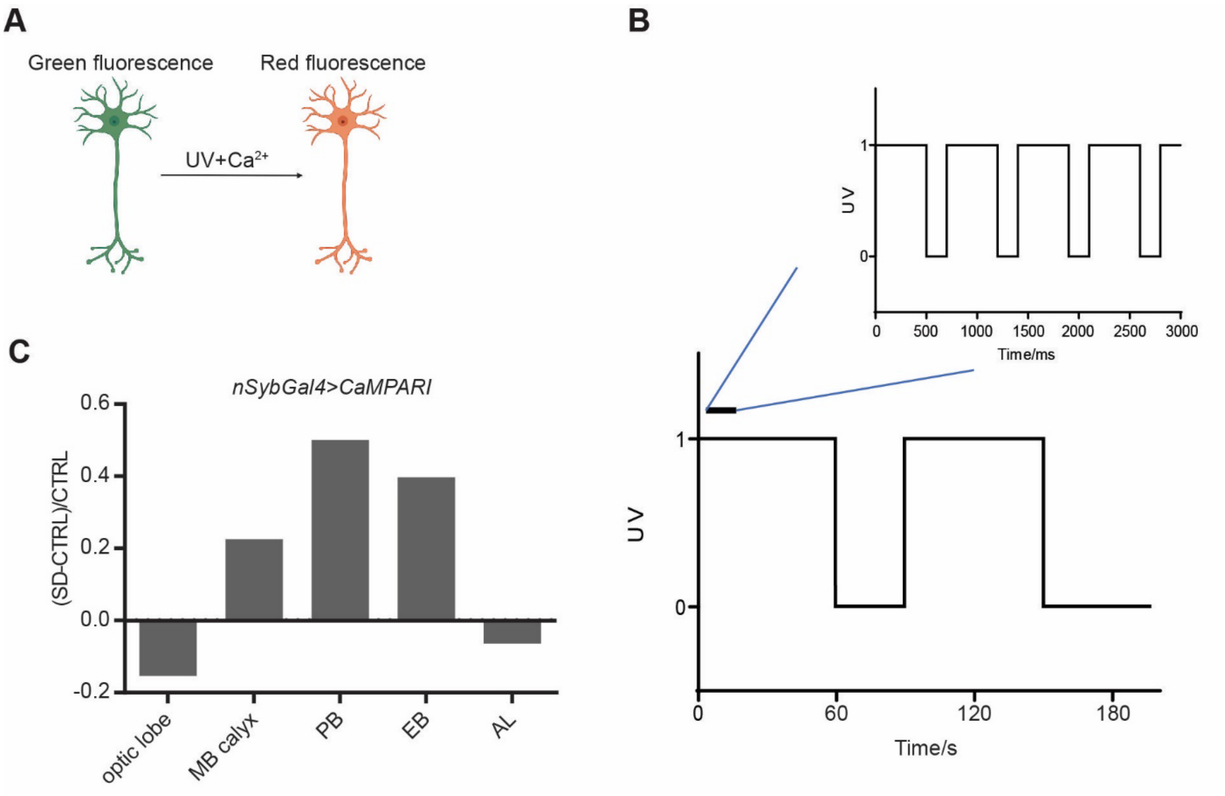


**Fig. S1. Calcium activity changes in different brain regions after sleep deprivation. (A)** A schematic representation of CaMPARI imaging. CaMPARI converts from green to red fluorescence in the presence of UV and high concentration of calcium. **(B)** Paradigm of UV pulsing in CaMPARI imaging. UV is pulsing with 500ms ON and 200ms OFF for 60 seconds followed by 30 seconds of break until a total duration of 150 seconds is reached. **(C)** Normalized change of CaMPARI $F_{red}/F_{green}$ ratio between the SD group after 13hrs sleep deprivation and the CTRL group at ZT1 in different brain regions.

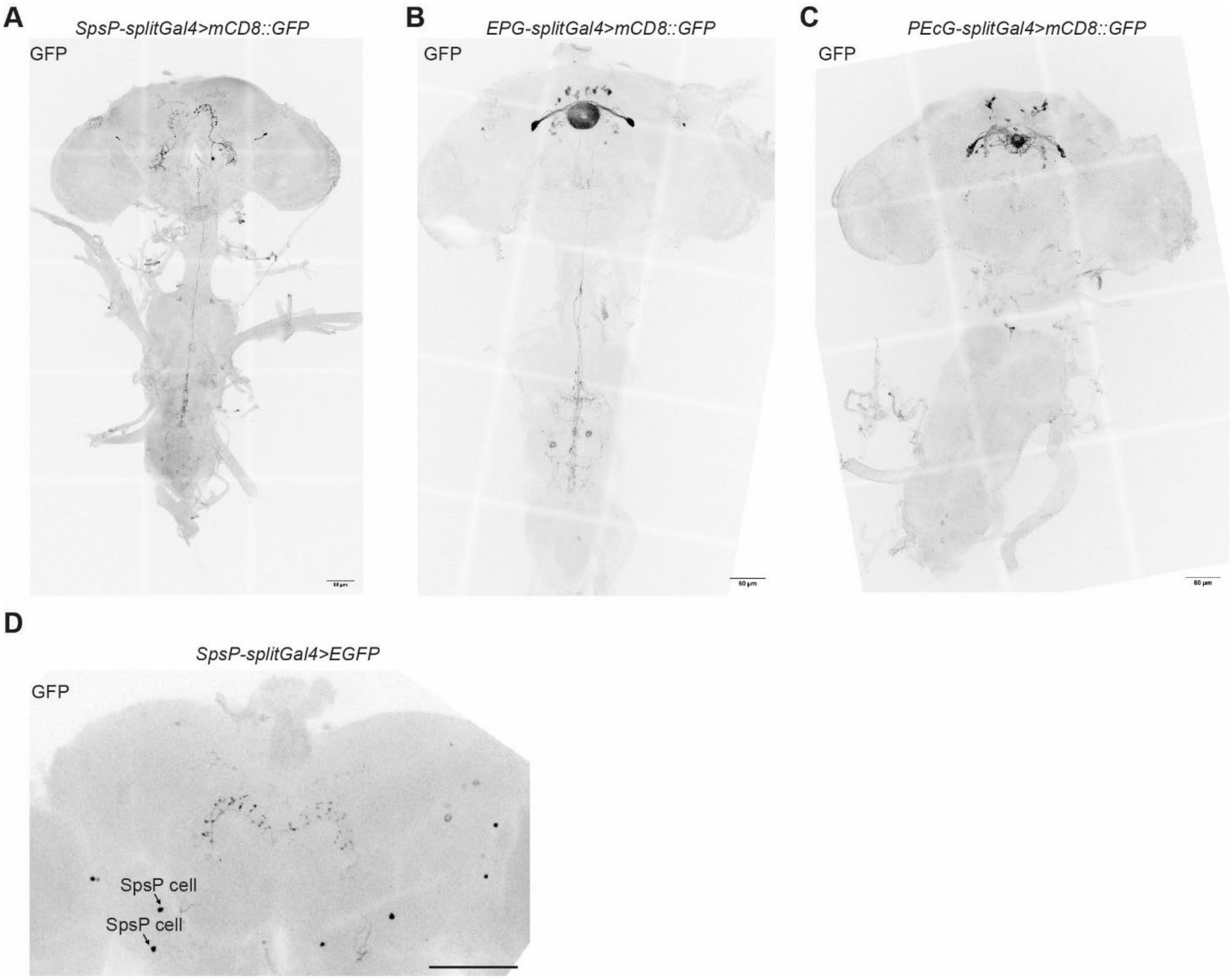


**Fig. S2. Candidate split drivers show sparse and specific expression patterns. (A-C)** Representative images of SpsP-splitGal4(ss52267)>mCD8::GFP **(A)**, EPG-splitGal4(ss50574)>mCD8::GFP **(B)**, and PEcG-splitGal4(ss02195)>mCD8::GFP (**C)** stained for GFP in the brain and the VNC. Scale bar: 60 µm. **(D)** Representative image of SpsP-splitGal4>EGFP stained for GFP in the brain. Arrows: Cell bodies of the SpsP cells in the left half-brain.  Scale bar: 60 µm.

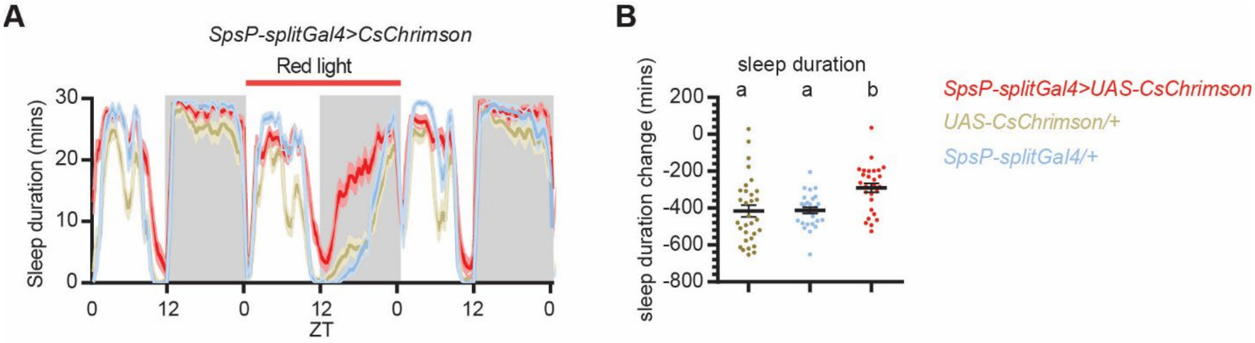


**Fig. S3. Optogenetic activation of SpsP neurons promote sleep.**

**(A-B)** Sleep profiles **(A)** and quantification of the changes of sleep duration **(B)** from ZT0-24 upon optogenetic activation of the SpsP neurons. Sleep profiles are averaged in 30 minute bins. Changes were calculated by subtracting the sleep duration of Day 1 ZT0-24 from that of Day 2. Shaded area/Error bars: S.E.M.. Red: SpsP-splitGal4>>UAS-CsChrimson (n=28); Yellow: UAS-CsChrimson/+ (n=31); Blue: SpsP-splitGal4/+ (n=29). Letters represent statistically distinct groups; $P < 0.01$, Kruskal–Wallis test followed by a post hoc Dunn's test. Male flies were used. Male flies were used.

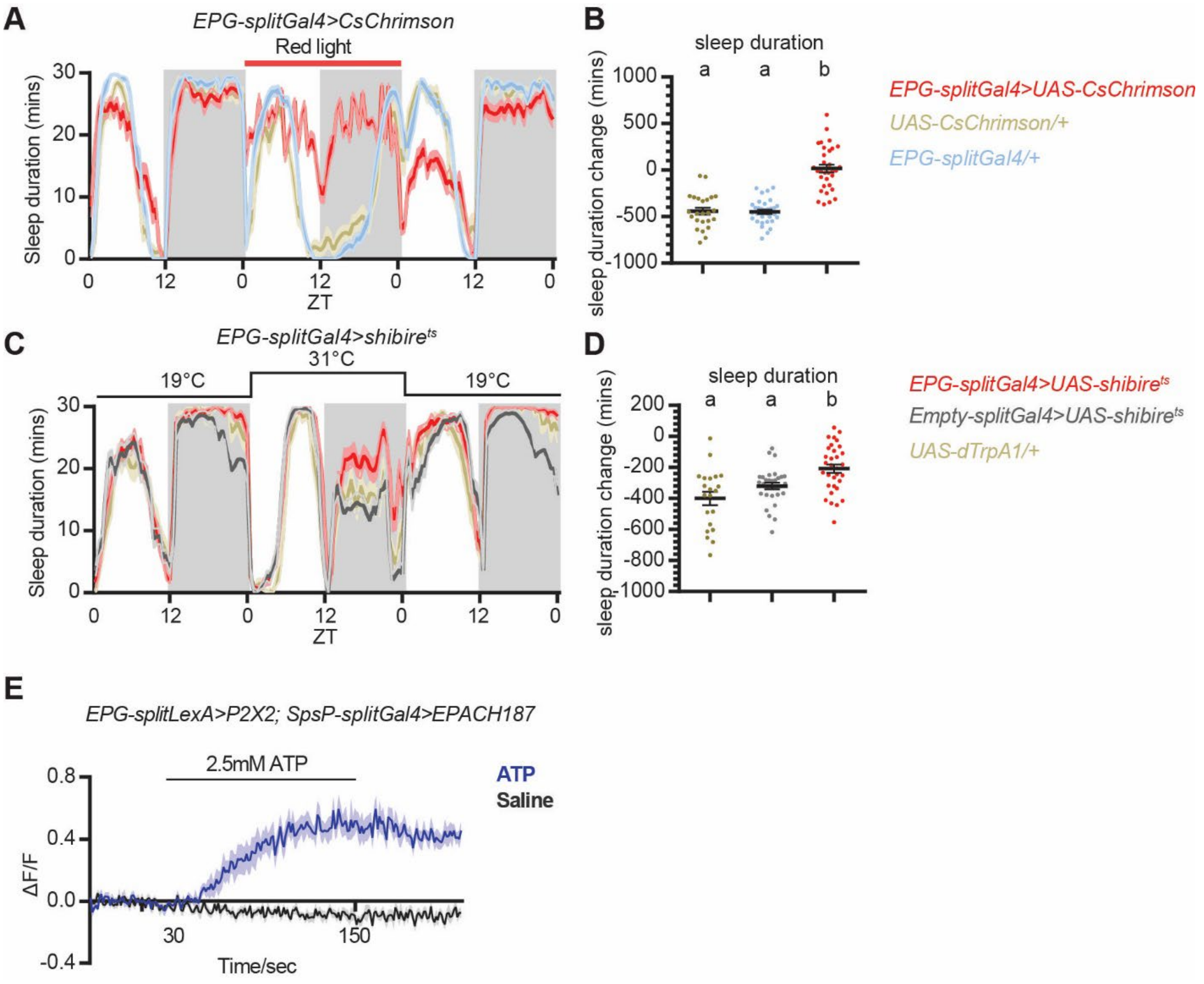


**Fig. S4. Activation of EPG neurons causes increased sleep and increased ATP level in SpsP neurons, whereas inhibition of E-PG neurons causes mildly increased nighttime sleep.** Sleep profiles **(A, C)** and quantification of the changes in sleep duration **(B, D)** from ZT0-24 upon optogenetic activation **(A)** or neurotransmitter release blocking **(C-D)** of the EPG neurons. Sleep profiles are averaged in 30 minute bins. Changes were calculated by subtracting the sleep duration of Day 1 ZT0-24 from that of Day 2. Shaded area/Error bars: S.E.M.. Red: EPG-splitGal4/UAS; Yellow: UAS/+; Blue: EPG-splitGal4/+; Gray: Empty-splitGal4/UAS. More than 20 male flies were used for each group. Letters represent statistically distinct groups; $P < 0.05$, Kruskal–Wallis test followed by a post hoc Dunn's test. **(E)** Average EPAC traces (ΔF/F:

inverse FRET signal) of SpsP cells in response to EPG activation. More than 7 male flies were used for each group. Error bars: S.E.M..

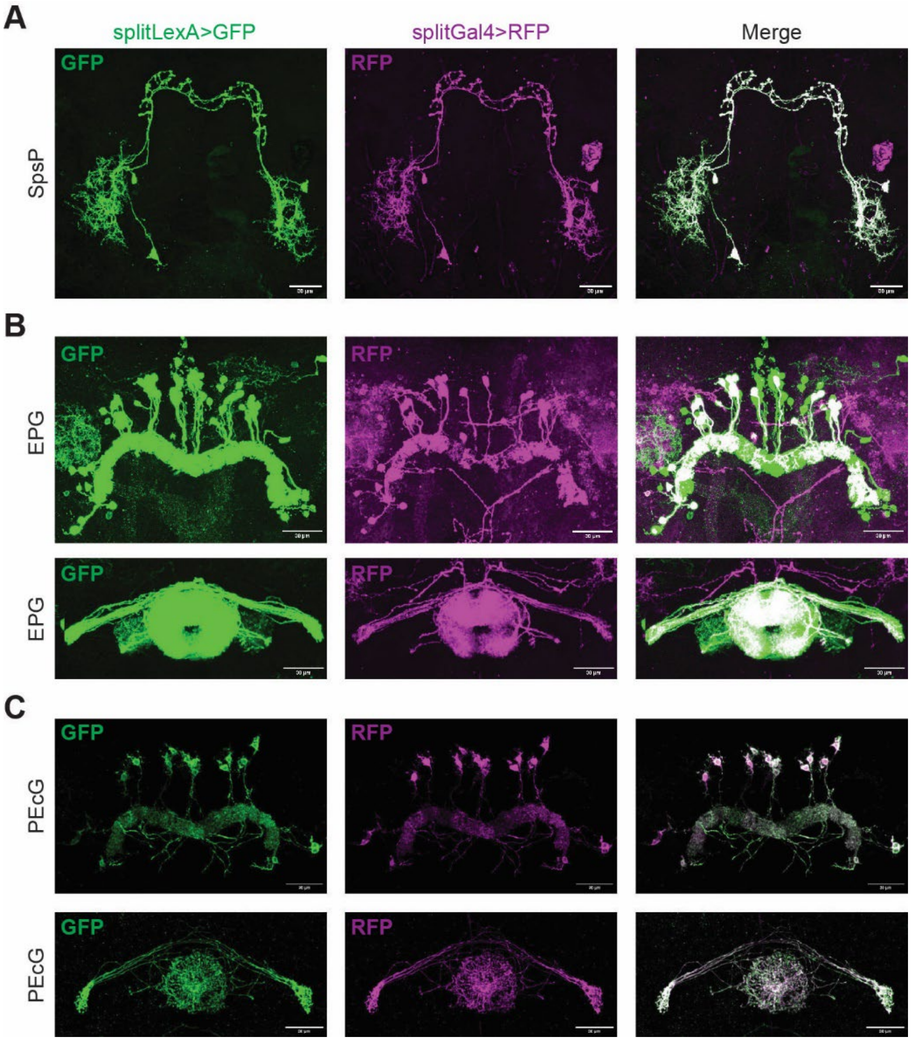


**Fig. S5. The expression patterns of split-LexAs recapitulate those of corresponding split-Gal4s.** Representative images of neurons labeled by split-LexA drivers (left) and split-Gal4 drivers of SpsP **(A)**, EPG **(B)**, and PEcG **(C)** stained for GFP and dsRed. Scale bar: 30 µm.

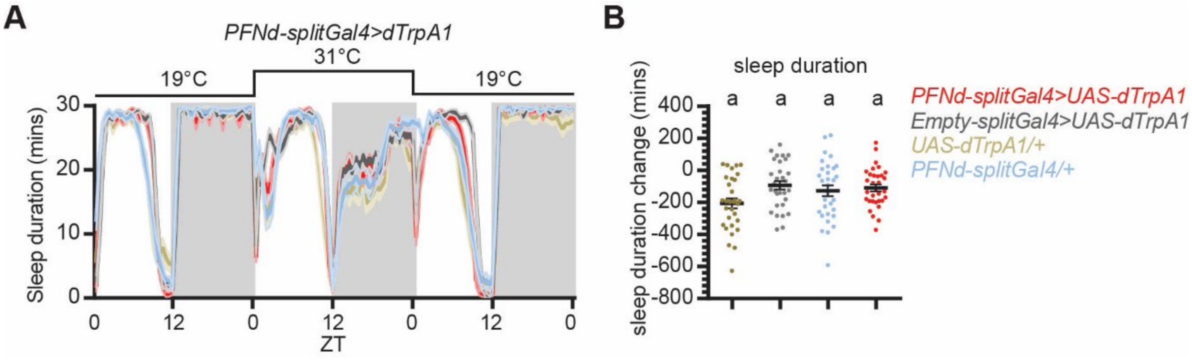


**Fig. S6. Activation of the PFNd neurons has no effect in sleep duration.** Sleep profiles **(A)** and quantification of the changes in sleep duration **(B)** from ZT0-24 upon thermogenetic activation of the PFNd neurons. Heatshock was from Day2 ZT0-24. Sleep profiles are averaged in 30 minute bins. Changes were calculated by subtracting the sleep duration of Day 1 ZT0-24 from that of Day 2. Shaded area/Error bars: S.E.M.. Red: PFNd-splitGal4>UAS-dTrpA1 (n=32); Gray: Empty-splitGal4/UAS-dTrpA1 (n=31); Yellow: UAS-dTrpA1/+ (n=31); Blue: PFNd-splitGal4/+ (n=31). Same letters represent statistically indistinct groups; P > 0.05, Kruskal–Wallis test followed by a post hoc Dunn’s test. Male flies were used.

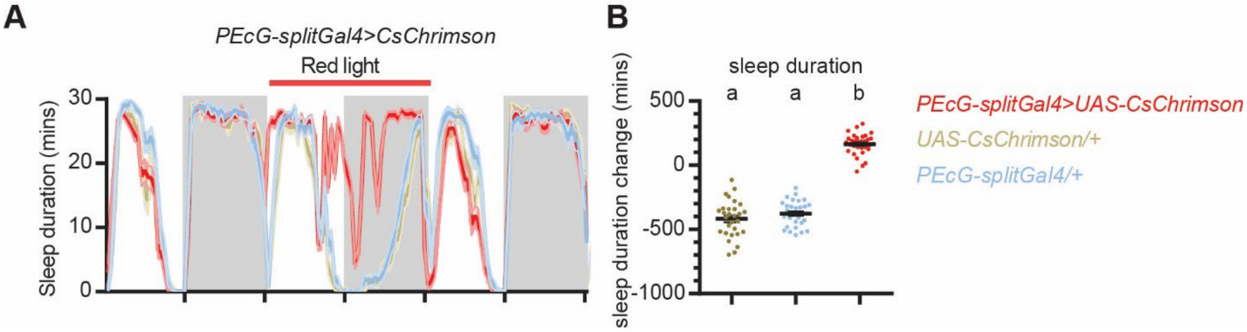


**Fig. S7. Activation of the PEcG neurons causes a sleep increase.** Sleep profiles **(A)** and quantification of the changes in sleep duration **(B)** of ZT0-24 upon optogenetic activation of the P-EcG neurons. Sleep profiles are averaged in 30 minute bins. Shaded area/Error bars: S.E.M.. Changes were calculated by subtracting the sleep duration of Day 1 ZT0-24 from that of Day 2. Red: P-EcG-splitGal4>UAS-CsChrimson (n=32); Yellow: UAS-CsChrimson/+ (n=29); Blue: P-EcG-splitGal4/+ (n=30). Letters represent statistically distinct groups; P < 0.0001, Kruskal–Wallis test followed by a post hoc Dunn’s test.

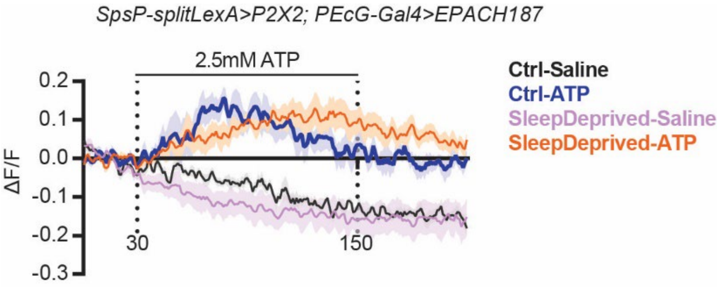


**Fig. S8. Activation of the SpsP neurons causes cAMP level increase in the P-EcG neurons.**

Averaged EPAC traces (ΔF/F, F: inverse FRET signal) of PEcG cells in response to ATP-induced SpsP activation. n=5 male flies for each group. Error bars: S.E.M..

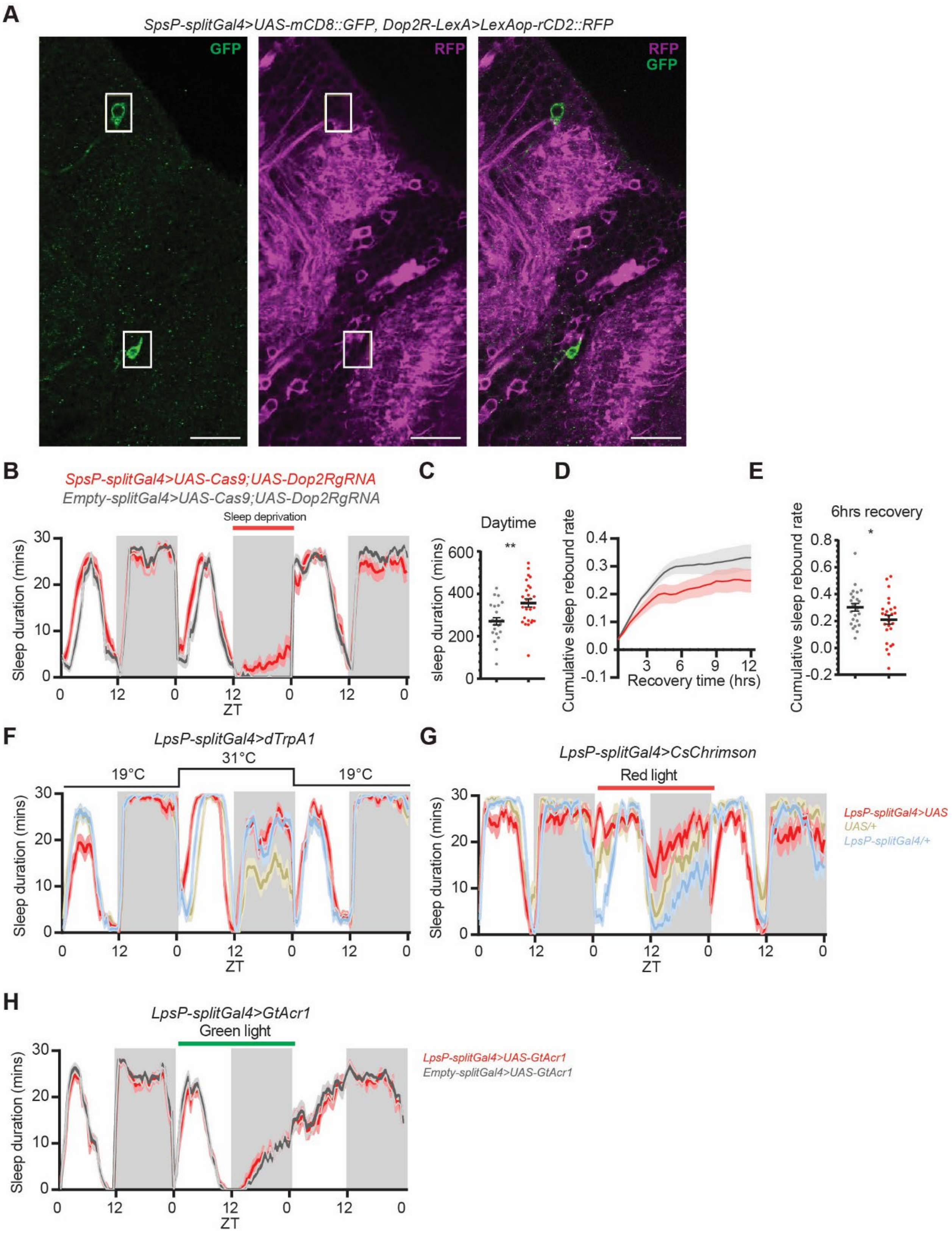


**Fig. S9. SpsP neurons integrate dopaminergic signaling to regulate sleep.**

**(A)** Representative images of the additional neurons labeled by the SpsP driver (green) and the Dop2R-expressing neurons (magenta) labeled by SpsP-splitGal4>UAS-mCD8::GFP, Dop2R-LexA>LexAop-rCD2::RFP stained for dsRed and GFP. Scale bar: 20 μm.

**(B-E)** Sleep profiles **(B)**, quantification of the daytime sleep duration from ZT0-12 **(C)**, cumulative sleep rebound rate curve in 12 hours **(D)**, and the quantification of cumulative sleep rebound rate after 6 hours of recovery sleep **(E)** in the experimental group and the control groups of SpsP specific Dop2R mutated flies. **(B)** and **(D)** are in 30 minute bins. Shaded area/Error bars: S.E.M.. Red: SpsP-splitGal4>UAS-Cas9; UAS-Dop2RgRNA (n=26); Gray: Empty-splitGal4>UAS-Cas9; UAS-Dop2RgRNA (n=24). *P<0.05, **P<0.01, Mann-Whitney test. Virgin female flies were used.

**(F-H)** Sleep profiles of thermogenetic activation **(F)**, optogenetic activation **(G)**, and optogenetic inhibition **(H)** of the dopaminergic LpsP neurons in 30 minute bins. Red: LpsP-splitGal4/UAS; Yellow: UAS/+; Blue: LpsP-splitGal4/+; Gray: Empty-splitGal4/UAS. More than 20 male flies were used for each group.
